# Engineered Secretory Immunoglobulin A provides insights on antibody-based effector mechanisms targeting *Clostridiodes difficile*

**DOI:** 10.1101/2023.11.08.566291

**Authors:** Sonya Kumar Bharathkar, Michael J. Miller, Beth M. Stadtmueller

## Abstract

Secretory (S) Immunoglobin (Ig) A is the predominant mucosal antibody, which mediates host interactions with commensal and pathogenic microbes, including *Clostridioides difficile*. SIgA adopts a polymeric IgA structure that is bound by secretory component (SC). Despite significance, how SIgA supports diverse effector mechanisms is poorly characterized and SIgA-based therapies nonexistent. We engineered chimeric (c) SIgAs, in which we replaced SC domain D2 with a single domain antibody or a monomeric fluorescent protein, allowing us to investigate and enhance SIgA effector mechanisms. cSIgAs exhibited increased neutralization potency against *C. difficile* toxins, promoted bacterial clumping and cell rupture, and decreased cytotoxicity. cSIgA also allowed us to visualize and/or quantify *C. difficile* morphological changes and clumping events. Results reveal mechanisms by which SIgA combats *C. difficile* infection, demonstrate that cSIgA design can modulate these mechanisms, and demonstrate cSIgA’s adaptability to modifications that might target a broad range of antigens and effector mechanisms.

## INTRODUCTION

Vertebrate epithelial barriers, largely formed by mucosal epithelia, are protected by specialized mucosal immune responses orchestrated in Mucosa Associated Lymphoid Tissues (MALT)[1]. These responses balance tolerance to commonly encountered antigens with mechanisms that promote pathogen clearance, but when perturbed can result in pathogen expansion, dysbiosis and disease[2],[3]. Mucosal antibodies play a major role in maintaining this balance; in humans, secretory (S) immunoglobulin (Ig) A functions as the predominant mucosal antibody, accounting for grams of antibodies secreted into the gut alone each day[4].

In humans, IgA exists in monomeric (m), polymeric (p) and secretory (S) forms, each mediating distinct effector functions. The mIgA comprises two heavy-chains (HC) and two light-chains (LC), that form two Fragment antigen binding (Fab) regions and one Fragment crystallizable (Fc) region[5],[6]. mIgA is predominant in serum and can initiate pro-inflammatory responses mediated by Fc-receptors (FcRs), such as Antibody dependent cellular phagocytosis (ADCP), antibody dependent cellular cytotoxicity (ADCC), and antibody mediated immunomodulation[7],[8]. On the contrary, pIgA contains two to five mIgAs, which plasma cells link together with a protein called the called joining-chain (JC); however, the predominant form of pIgA in humans is dimeric (d). All pIgA can bind the polymeric Immunoglobin receptor (pIgR), which transcytoses the complex from the basolateral surface of epithelial cells to the apical cell surface. There the pIgR ectodomain is proteolytically cleaved, releasing SIgA, a complex of the pIgA and the cleaved pIgR ectodomain (secretory component; SC), into mucosal secretions[9],[10].

In contrast to mIgA, SIgA effector mechanisms are largely anti-inflammatory and balance the removal of pathogens with tolerance for commensal microorganisms[11]. This delicate balance is accomplished through a combination of poorly understood innate-like and adaptive mechanisms[10],[12]. Adaptive mechanisms are mediated by Fabs, which can form high-affinity interactions with antigens on microbial surfaces or extracellular pathogenic secretions such as toxins. A subset of antigen-SIgA complexes can potentially interact with host receptors and may contribute to proinflammatory responses[13],[14],[15]. The majority of SIgA interactions result in SIgA coating or cross-linking of microbes, which typically results in immune exclusion, a mechanism that prevents the breach of epithelial barriers by microbes or toxins and promotes their removal via peristalsis[16]. On a molecular level, SIgA can mediate crosslinking of multiple antigens (e.g., surface proteins or carbohydrates) on one microbe, or crosslinking of identical antigens on two microbes. Microbial crosslinking is presumed to be supported by SIgA’s polymeric structure and can occur through at least two distinct mechanisms, classical agglutination, or enchained growth. Classical agglutination results from collisions between SIgA-coated microbes, which cause them to stick together, and is favored when the local microbial concentrations and associated collisions are high. Enchained-growth results from SIgA-coated mother cells dividing and resulting in daughter cells remaining linked. In enchained-growth, successive divisions, rather than collisions, can result in clumps of crosslinked microbes. Agglutination and enchained growth are not mutually exclusive and depend on many factors including microbial morphology and concentration, antigen expression, and distribution, as well as antibody affinity and other factors such as location and concentration[16],[17]. SIgA’s innate-like mechanisms are thought to be mediated primarily through non-specific interactions with the constitutive parts of SIgA, like HC, LC, JC, SC, and the N and O linked glycans, which may provide passive immunity through non-specific interactions that can also lead to immune exclusion[12],[18],[19]. Together, SIgA’s innate and adaptive mechanisms can limit microbial adherence to host epithelial cells, promote pathogen clearance, or promote colonization of some commensal species. It is also notable that SIgA binding to microbes can impact the microbe in many ways, potentially altering gene expression, division, and other physiological processes[16],[17]. All of these processes are supported by SIgA’s unique molecular structure, which is characterized by an asymmetric arrangement of complex components that constrains the positions of the Fabs and leaves two FcR binding sites and the SC sterically accessible[17]. However, SIgA’s structure-function relationships remain poorly understood.

Deciphering the mechanisms behind SIgA-antigen interactions is critical to understanding and treating a vast array of diseases, yet the diversity of effector mechanisms and outcomes implies that understanding many antigen-specific processes is necessary to achieve this. Of particular interest are SIgA interactions with human gut pathogen *Clostridiodes difficile*, which causes *Clostridiodes difficile* Infection (CDI). CDI is one of the most prevalent nosocomial infections in first world countries[20]. Typically, the disease damages epithelial barriers leading to pseudomembranous colitis, that result is 2.7% of CDI related death upon primary infection, and 25.4% of CDI related death upon recurrence. Additionally, 35% of CDI recovered patients develop recurrence of the disease, and 65% of those who experience one recurrence go on to have yet another recurrence[21]. The barrier damage is caused primarily by *Clostridiodes difficile* toxins, TcdA and TcdB, which are the main virulence factors and are cytotoxic to host cells[22], but the disease is exacerbated by rapid growth of *C. difficile,* and the incomplete pathogen clearance upon treatment[20]. Standard CDI treatments, including metronidazole, fidaxomicin and vancomycin treatment often fail to completely clear the pathogen and target commensal species leading to prolonged gut dysbiosis and recurrence; and they also fail to target *C. difficile* toxins, TcdA and TcdB, and *C. difficile* spores [23],[24], both of which contribute to recurrence [25]. Recently, the monoclonal antibody (mAb) Bezlotoxumab, which targets TcdB, became the first FDA approved IgG1 mAb that can limit TcdB cytotoxicity when delivered intravenously along with antibiotic treatment[26],[27]. However, Bezlotoxumab cannot limit *C. difficile* growth and based on the intravenous administration approach[26], likely targets internalized toxins, presumably after damage has already occurred rather than neutralizing or excluding the toxin on the apical side of intestinal epithelial cells. Human hosts develop SIgA targeting *C. difficile* antigens[28], implying that SIgA offers natural protection against CDI and may hold therapeutic potential. Yet, the SIgA dependent effector mechanisms that promote successful *C. difficile* clearance or limit toxin cytotoxicity remain poorly understood.

In sum, SIgA is effectively an antibody multi-tool; yet knowledge on how each part of the tool works against diverse mucosal antigens and how it can be harnessed to benefit health remains limited. Here we address this challenge by using structure-based engineering to create a series of chimeric SIgA with novel bifunctional modalities that confer either (1) enhanced antigen binding through an introduced single domain antibody (sdAb) or (2) enhanced detection through an introduced monomeric fluorescent protein (mFP). Using a library of natural and chimeric (c) SC and SIgAs, we investigated SIgA-dependent neutralization of *C. difficile* and its toxins and evaluated the potential of cSC and cSIgA to enhance neutralization. Our results provide mechanistic insights on SIgA-dependent effector functions, demonstrate an advantage of SIgA over monomeric counterparts and remarkably, reveal that this can be further enhanced through SC-based engineering. Broadly, these findings shed new light on SIgA effector mechanisms applicable to diverse antigens and offer opportunities to develop cSIgA tools for investigating and treating disease.

## RESULTS

The SC and SIgA structures revealed that the majority of SC D2 and its Complementarity Determining Region (CDR)-like loops are solvent accessible in both unliganded and liganded forms and do not directly contribute to binding pIgA [29],[30],[31],[32], making D2 an attractive target for engineering bifunctional modalities into SC and SIgA (Figure 1A). To explore this possibility, we created expression constructs, in which D2 was replaced with a sdAb or a mFP (Figure 1B). Using these strategies, we made a library of chimeric SC (cSC), each containing a sdAb that binds a unique antigen, or containing a mFP. Additionally, we co-expressed cSC with JC and IgA (or Fcα-fused-sdAb) variants to generate a library of chimeric SIgA (cSIgA), in which the cSC recognized an antigen or conferred fluorescence, and the Fab (or fused sdAb) recognized either the same antigen or a different antigen. These strategies preserved the conformational advantages (e.g. bend, tilt, avidity) of SIgA while providing additional, modular functions and allowed us to use cSIgA as a versatile tool to study the neutralization potential and effects of cSIgA on its targets based on customized modifications.

**Figure 1:**
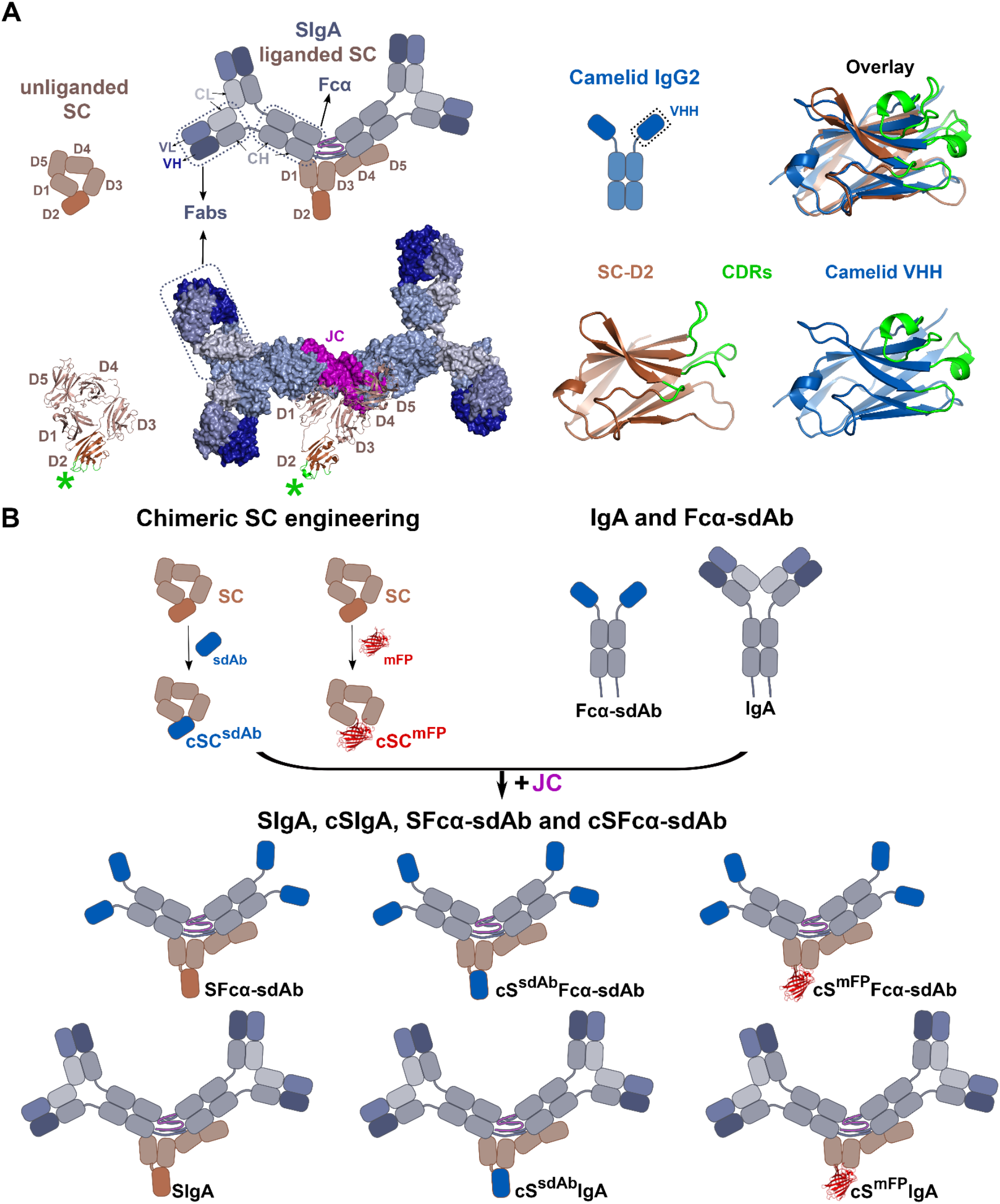
Chimeric SC and SIgA engineering strategy. (A) The schematic representations of SC, SIgA and camelid IgG2, and structural representation of SC (PDB: 5D4K) and SIgA (PDB: 7JG2). The SC has five Ig-like domains (D1-D5; brown) which exist in a closed unliganded conformation and open SIgA-liganded conformation. The D2 (dark brown) CDR-like loops (green) are solvent accessible in both conformations. The SIgA is composed of one SC, one JC and of two IgA monomers resulting in two pairs of Fabs and two Fcs (indicated). The SC-D2 (brown) and Camelid VHH (blue) structure and their CDRs (green) are shown, along with the structures aligned, showing CDRs oriented in the same direction. (B) Schematics of cSC, in which sdAbs or mFPs replace D2, are combined with JC and IgA or Fcα-fused-sdAb, to create cSIgA variants.

### The SC D2 can be replaced with sdAbs

With a goal of targeting CDI virulence factors, we generated cSC and cSIgA variants that incorporated sdAbs recognizing the *C. difficile* large clostridial toxins (LCT), TcdA and TcdB, or recognizing other antigens. The resulting cSC exhibited high tolerance for a wide range of sdAb, and VH and VL domains, as evidenced by monodisperse SEC elution profiles for the resulting cSCs (Figure S1A). To test the ability of cSCs to bind target antigens, we focused efforts on a cSC variant, in which the D2 was replaced with sdAb, sdA20.1, called cSC^A20.1^. The sdA20.1 binds the TcdA combined repetitive oligopeptide (CROP) domain, (Figure 2A), and is thought to block interactions between putative CROP receptor binding motifs and host-epithelial cell receptors, thereby preventing toxin internalization[33],[34].

**Figure 2:**
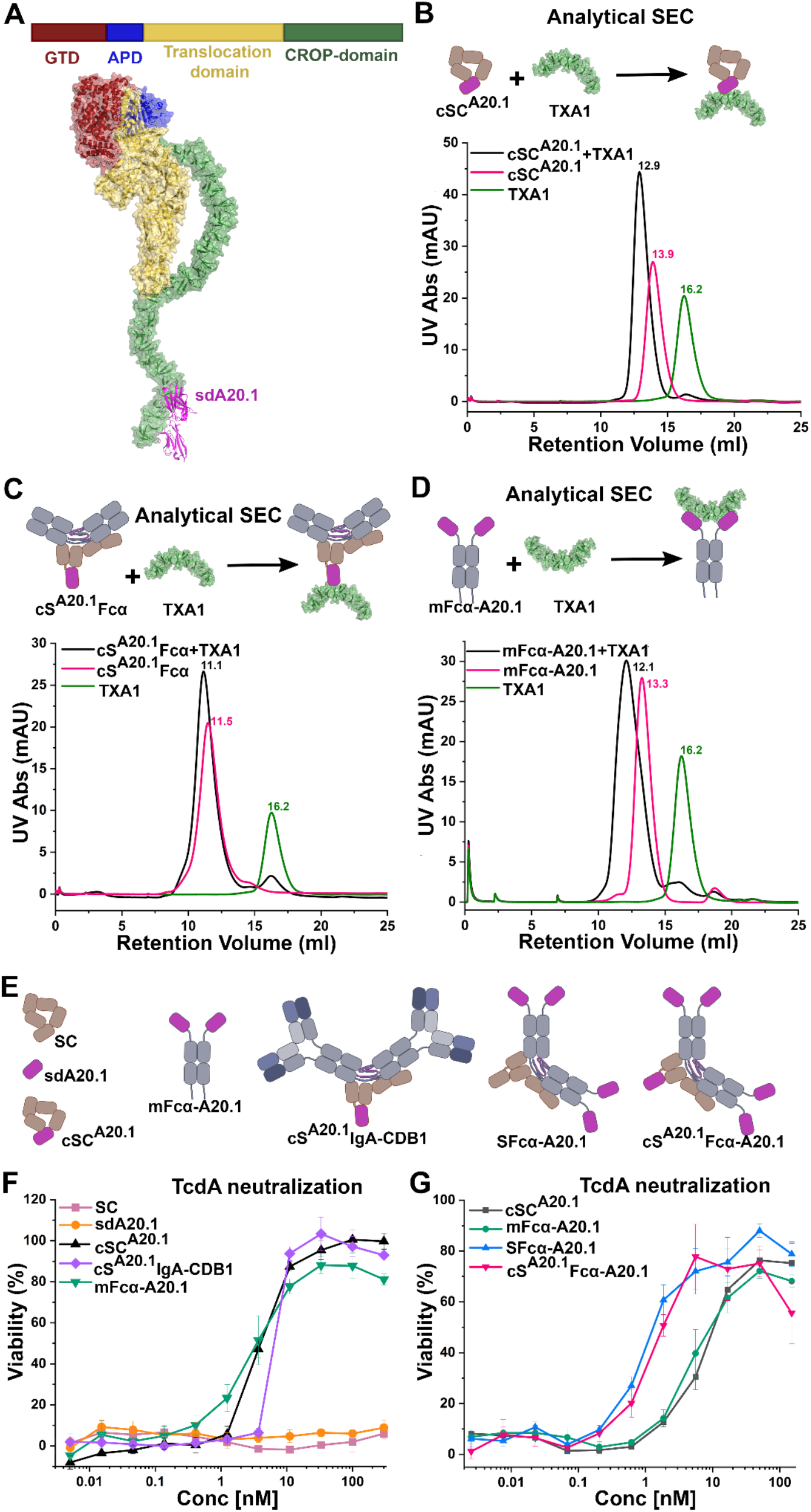
**cSC and cSIgA bind and neutralize TcdA. (**A) Schematic of TcdA domain organization (top) and TcdA-sdA20.1 structure (bottom; combined structures with PDB code:7U1Z and 4NBY). The TcdA domains are colored the same in the schematic and the structure, and sdA20.1 is colored magenta. (B) Schematic and analytical SEC elution profiles of cSC^A20.1^, TcdA fragment TXA1, and the cSC^A20.1^-TXA1 complex. (C) Schematic and SEC elution profiles of cS^A20.1^Fcα, TXA1, and the cS^A20.1^Fcα-TXA1 complex. (D) Schematic and SEC elution profiles of mFcα-A20.1, TXA1, and a mFcα-A20.1-TXA1 complex. The analytical SEC was carried out by incubation of equimolar amounts of TXA1 and cSC^A20.1^, cS^A20.1^Fcα or mFcα-A20.1 at room temperature for 1hr, followed by chilling the mixture to 4 degrees and then carrying out SEC. Similarly, TXA1, cSC^A20.1^, cS^A20.1^Fcα and mFcα-A20.1 were individually subjected to SEC for comparison. (E) Schematic of antibodies used for TcdA neutralization assays. (F) Neutralization profiles of the indicated molecules against 50pM TcdA, where cSC^A20.1^ and cS^A20.1^IgA-CDB1 are the chimeras compared against the neutralization capability of sdA20.1, and mFcα-A20.1. (G) The concentration dependent neutralization of 50pM TcdA, where potential for increasing the number of binding sites for TcdA neutralization was tested with different chimeras involving different number of sdA20.1 moieties. *See also Figure S1*.

Specifically, we produced cSC^A20.1^ (unliganded) and cS^A20.1^Fcα (liganded), a cSIgA lacking Fabs, and tested their binding to TcdA-CROP fragment, TXA1, using analytical SEC (Figure 2B and 2C; Figure S1C). Results revealed peaks corresponding to TXA1-cSC^A20.1^ and TXA1-cS^A20.1^Fcα complexes with lower retention volume than controls, indicating that cSC^A20.1^ can bind the TXA1 antigen in both its unliganded and liganded conformations. The corresponding sdAb called sdA20.1 and monomeric Fcα-fused variant, called mFcα-A20.1, also showed *in-vitro* binding that was verified by Analytical SEC as controls (Figure 2D; Figure S1B-C). Together, results demonstrate that the replacement of SC-D2 with sdAbs is an effective means of adding antigen binding capacity to SC and SIgA.

### cSC and cSIgA can neutralize Toxins

Having developed a means to add a sdAb and its binding specificity to cSC and cSFcα, we aimed to test the neutralization potency of: (1) unliganded and liganded versions cSC^A20.1^ and (2) natural and chimeric variants containing different numbers of antigen binding sites (provided by sdAbs or Fabs); for example, one site on cSC, two sites on mIgA, four or more sites on SIgA, and five or more sites on cSIgA. In the case of SIgA and cSIgA, antigen binding specificity can be provided by cSC, Fabs, or sdAbs linked to Fcα. While Fabs provide typical mammalian antigen binding sites, we reasoned that fusing sdAbs to Fcα would allow us to test how increasing the number of identical antigen binding sites (e.g., the number of sdA20.1) in chimeric variants would influence neutralization potency. Accordingly, we created mFcα-A20.1, which fuses Fcα with sdA20.1, (the same sdAb included in cSC^A20.1^), and confirmed binding of mFcα-A20.1 to TXA1 using analytical SEC (Figure 2D; FigureS1C). Additionally, we produced, sdA20.1, cSC^A20.1^, mFcα-A20.1, the secretory form-SFcα-A20.1, and chimeric Secretory form-cS^A20.1^Fcα-A20.1, as well as a chimeric SIgA, cS^A20.1^IgA CDB1 (Figure 2E). The Fabs in cS^A20.1^IgA-CDB1, CDB1-Fab[35], do not bind to TcdA and therefore provide a means to test the potency of cSC^A20.1^ in its liganded conformation with Fabs present.

To determine the neutralization potency of chimeric antibody variants targeting TcdA, we conducted Vero cell-based cytotoxicity neutralization assays, in which Vero cell cultures were incubated with varying concentrations of antibodies and constant concentration of 50pM TcdA, and their cell viability was measured after ∼65hrs by an Alamar Blue assay[36]. Cell viability greater than 50%, is indicative of toxin neutralization; 50pM TcdA alone that causes ∼100% Vero-cell death in the absence of antibodies.

We initially carried out cytotoxicity neutralization assays comparing cSC^A20.1^ (unliganded cSC), cS^A20.1^IgA-CDB1 (liganded cSC), mFcα-A20.1, sdA20.1, and SC (non-binding control). Resulting neutralization curves demonstrated that cSC^A20.1^ and cS^A20.1^IgA-CDB1 can readily neutralize the cytotoxic effects of TcdA whereas the wild-type SC cannot (Figure 2F). Notably, we observed similar levels of neutralization for unliganded and liganded cSC^A20.1^ and mFcα-A20.1; however, sdA20.1 on its own failed to neutralize TcdA, despite evidence for toxin fragment binding(Figure 2F)[34]. It is notable that chimeras containing cSC^A20.1^ (one copy of A20.1) performed similarly to mFcα-A20.1 (two copies of A20.1), whereas sdA20.1 on its own failed to neutralize. This suggests that sterically bulkier A20.1 variants, such as cSC^A20.1^, cS^A20.1^Fcα, and mFcα-A20.1, are better at neutralizing the TcdA, perhaps because they better shield TcdA interactions with host cell receptors (Figure 2F). Similar experiments were carried out with cSC^5D^, where sd5D binds to TcdB translocation domain[37], in both liganded and unliganded forms, cSC^5D^ showed neutralization capability for TcdB (Figure S1D and S1E).

Subsequently, we carried out cytotoxicity neutralization assays comparing SFcα-A20.1 (four or more copies of sdA20.1) and cS^A20.1^FcαA20.1 (five or more copies of sdA20.1). SFcα-A20.1 includes a predominantly dimeric Fcα comprising four heavy chains, each fused to one sdA20.1, implying that SFcα-A20.1 contains four copies of sdA20.1 and cS^A20.1^FcαA20.1 contains five copies of sdA20.1. However, like naturally produced SIgA, a minor percentage of SFcα molecules may adopt larger polymeric states (e.g. tetramers and pentamers). These chimeras exhibited increased neutralization potency compared to variants containing one to two copies of sdA20.1, indicating that for a chimeric SIgA-based antibody targeting TcdA, increasing the number of sdAb can be functionally advantageous (Figure 2G). Together data indicate that one copy of sdA20.1 in cSC provides similar protection to two copies of sdA20.1 in mFcα, both of which provide a considerable advantage over sdA20.1 alone. Neutralization potency could be further improved when four or more copies of sdA20.1 were included in the form of SFcα or cSFcα antibodies.

### cSIgA targeting multiple antigens show synergistic effects on TcdA and TcdB neutralization

Given similar neutralization potencies of cSC^A20.1^ and mFcα-A20.1, we reasoned that we could increase the neutralization breadth and/or potency of cSIgAs by creating bispecific cSIgAs targeting multiple domains on one toxin and/or targeting two different toxins. For this purpose we sought a TcdB-targeting Fab to combine with cSC^A20.1,^ and therefore selected PA41, which is reported to neutralize TcdB by binding the TcdB’s Glucosyl Transferase Domain(GTD); the main enzymatic portion of the toxin (Figure 3A)[38]. Neutralization assays containing either TcdA or TcdB revealed that mIgA-PA41 neutralized both TcdB (its known antigen) and, to a lesser extent, TcdA (Figure S2A). The unexpected PA41-dependent neutralization of TcdA may result from sequence similarity in the TcdA and TcdB GTD domains and the PA41 binding sites(Figure S2B).

**Figure 3:**
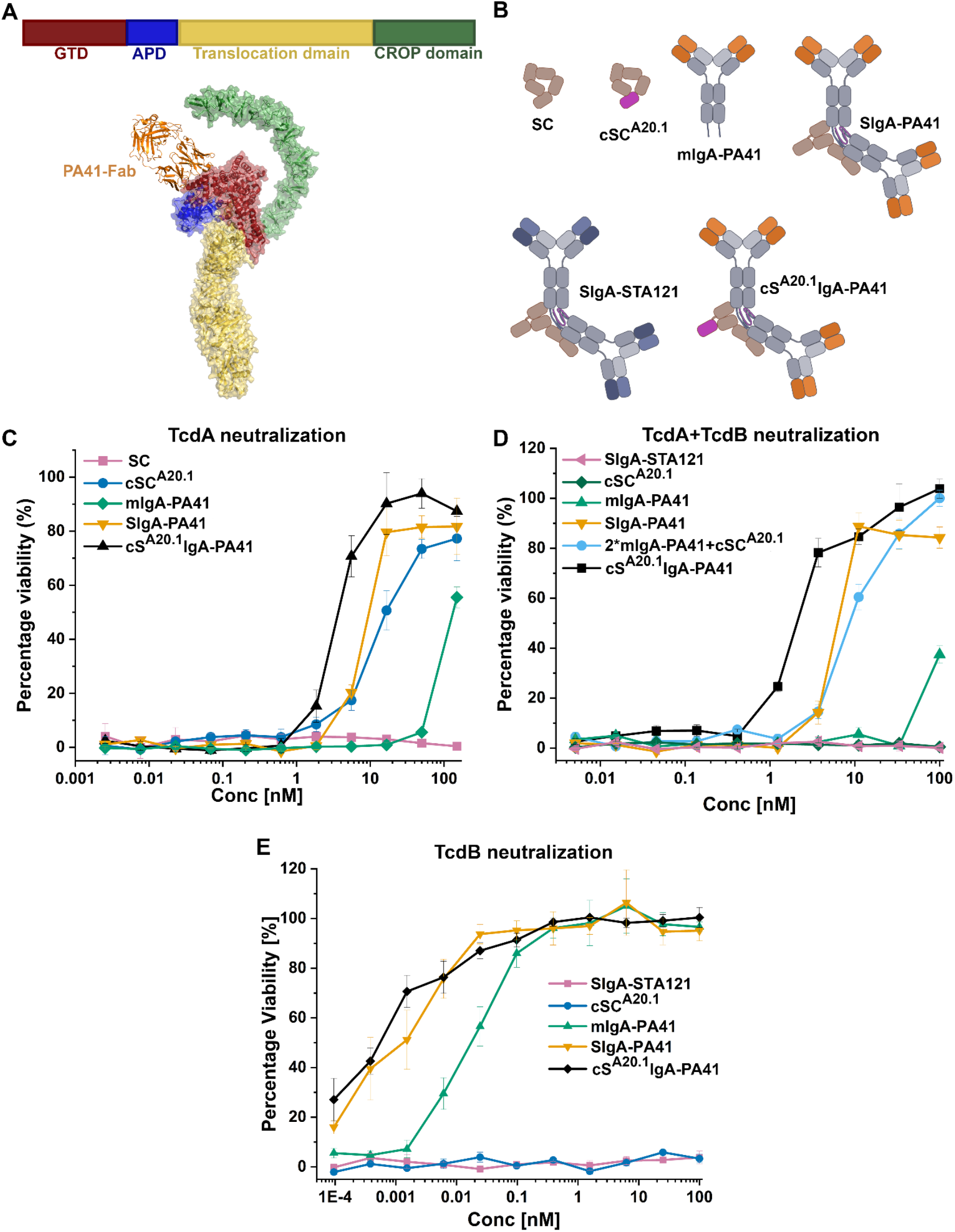
**cSIgA targeting multiple antigens show synergistic effects on TcdA and TcdB neutralization. (**A) Shows the domain organization of TcdB (top) and the binding site of PA41-Fab on the GTD domain of TcdB (PDB: 5VQM and 6OQ5). (B) Schematic representation of antibodies used in cytotoxicity assays including SC, cSC^A20.1^, mIgA-PA41, SIgA-PA41, SIgA-STA121 and cS^A20.1^IgA-PA41. (C) Concentration dependent neutralization of TcdA (50pM) by the indicated molecules including the bispecific antibody cS^A20.1^IgA-PA41. (D) Concentration dependent neutralization of TcdA (50pM) and TcdB (4pM) by the indicated molecules, including the bispecific antibody cS^A20.1^IgA-PA41. (E) Concentration dependent neutralization of TcdB (4pM) by the indicated molecules, including the bispecific antibody cS^A20.1^IgA-PA41. *See also Figure S2 and Figure S3*.

Accordingly, we created mIgA-PA41, SIgA-PA41 and cS^A20.1^IgA-PA41 and control antibodies (Figure 3B) and assayed neutralization potency against TcdA (50pM), TcdB (4pM) or TcdA (50pM) + TcdB (4pM) in Vero-cell based cytotoxicity assays. The inclusion of PA41 and A20.1 in a single molecule tests the neutralization potency of bi-specific cS^A20.1^IgA-PA41 against two different epitopes on the TcdA antigen (GTD and CROPs) and one epitope on the TcdB (GTD).

Notably, in assays containing only TcdA, neutralization curves revealed that cS^A20.1^IgA-PA41 has enhanced TcdA (50pM) neutralization potency compared to chimeric molecules that incorporate cSC^A20.1^ or PA41 alone (Figure 3C). Subsequently, we tested the neutralization potency of cS^A20.1^IgA-PA41 in Vero cell cytotoxicity assays containing both TcdA and TcdB, which also revealed enhanced neutralization of TcdA (50pM) and TcdB (4pM) compared to proteins and complexes that incorporate cSC^A20.1^ or PA41 alone (Figure 3D). Additionally, the cS^A20.1^IgA-PA41 and SIgA-PA41 revealed similar neutralization potency against TcdB (4pM), likely because cSC^A20.1^ does not neutralize TcdB, rendering the additional binding site insignificant (Figure 3E).

Similar experiments were carried out with the bispecific antibody cS^5D^IgA-Pa41 against TcdA (50pM), TcdB (4pM) and TcdA(50pM) + TcdB (4pM). Here, cS^5D^IgA-Pa41 shows neutralization potency against two epitopes of TcdB (GTD and translocation domain) and one epitope of TcdA (GTD). In these cases, cS^5D^IgA-PA4 showed enhanced neutralization in all cases (Figure S3A to S3D).

Together, these findings indicate that utilizing cSC with IgA Fabs to merge two different epitope binding specificities into a single bi-specific cSIgA can result in favorable synergistic effects that enhance neutralization. In the context of antibodies targeting *C. difficile* toxins, these results imply that the use of the PA41-Fab in a cSIgA could provide increased breath against multiple toxins and increased potency against one toxin.

### cSC facilitates visualization of SIgA targeting the *C. difficile*

Having demonstrated the neutralization potential of cSC and cSIgA against soluble *C. difficile* antigens we sought to investigate SIgA interactions with *C. difficile* surface antigens. As noted, SIgA effector mechanisms are diverse and are dependent on numerous variables, and therefore understanding how SIgAs affect different stages of pathogenesis is critical for understanding and treating CDI and other mucosal diseases. To accomplish this, we focused on antibodies that target the *C. difficile* Surface layer protein (SLP; gene *SlpA*), which forms the outermost layer of the bacterium and plays a major role in host cell adherence (Figure 4) [39],[40],[41]. The mature SLP is composed of a high molecular weight (HMW) and low molecular weight (LMW) SLP subunits linked non-covalently to each other and assembled as a paracrystalline array on the surface of bacterium above the cell-wall (Figure 4)[40] The HMW-SLP is highly conserved across different *C. difficile* strains while LMW-SLP exhibit variance in different serotypes of *C. difficile*[42],[44]. Evidence suggests that human hosts develop immune responses against *C. difficile* SLP[28],[42],[45]; yet, antibody specificity and how they influence CDI remains poorly understood[39]. For example, SIgA could coat individual bacteria and/or crosslink multiple copies of bacteria either through enchained growth or classical agglutination, mechanisms which have the potential to limit bacterial growth, limit adherence to host cells and/or effect other downstream functions. To investigate this, we chose sdAb, sdVHH5 (alias sdCD5SLP), which is reported to bind the low-molecular weight portion of SLP (LMW-SLP)[46]. We incorporated sdCD5SLP into cSC as cSC^CD5SLP^ in liganded and unliganded versions, and Fcα-fusion as mFcα-CD5SLP. We verified antigen binding of sdCD5SLP, cSC^CD5SLP^ (unliganded), cS^CD5SLP^Fcα (liganded) and mFcα-CD5SLP with recombinant LMW-SLP, by analytical SEC (Figure 4B and 4C; Figure S4B to S4E).

**Figure 4:**
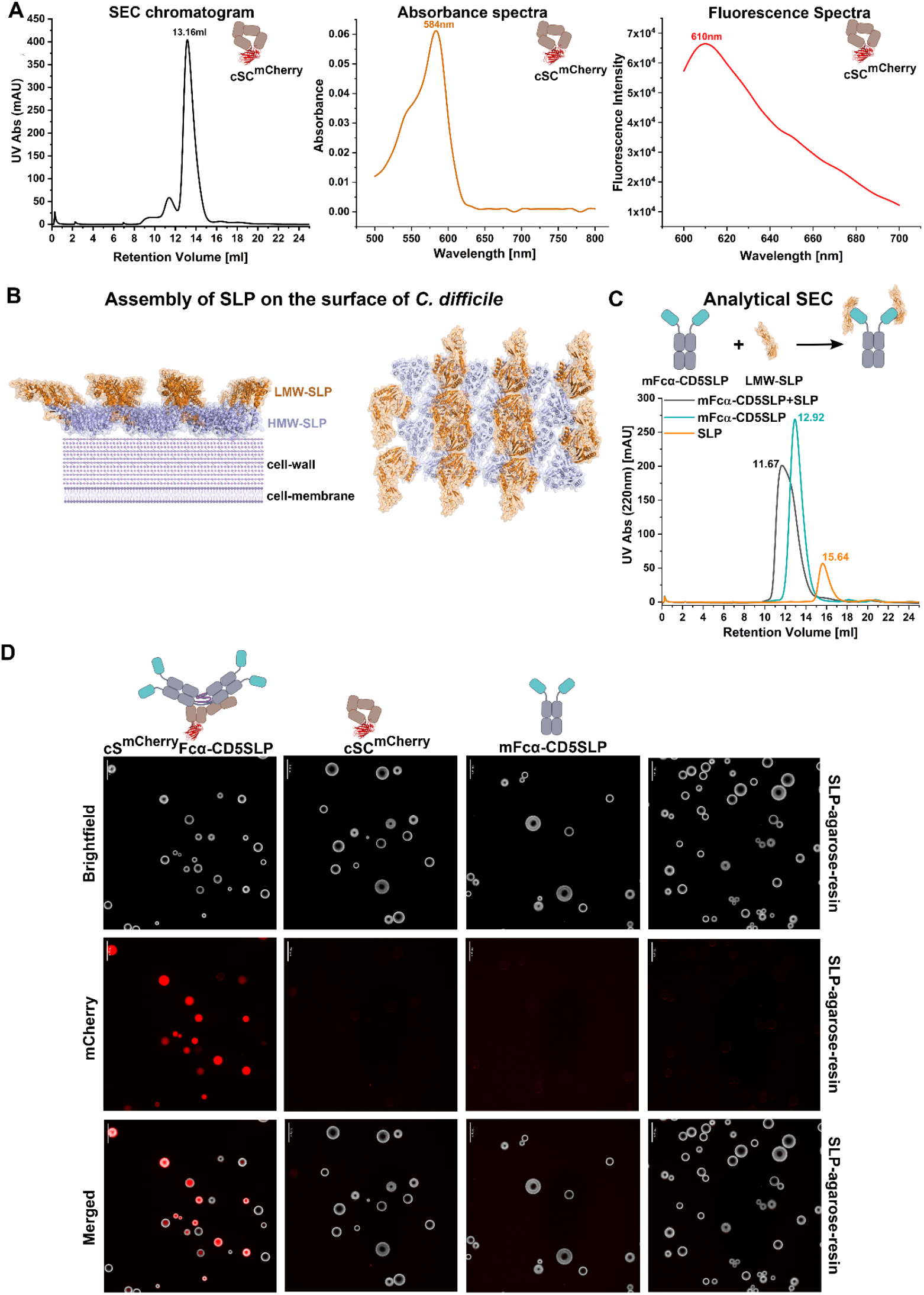
SC-D2 tolerates domain replacement with mFPs and can be used to make polymeric fluorescent antibodies for antigen specific imaging. (A) The SEC chromatogram of cSC^mCherry^, showing a monodisperse peak, followed by the absorption spectra of 1μM cSC^mCherry^, and fluorescence spectra of cSC^mCherry^ upon excitation with 560nm. (B) Schematic showing the assembly of surface layer proteins after the SlpA is cleaved to form high molecular weight (HMW) and Low molecular weight (LMW) fragments, the latter of which forms the outermost surface of the bacteria and is solvent accessible (PDB: 7QGQ). (C) Analytical SEC showing the binding of mFcα-CD5SLP binding to LMW-SLP. (D) Brightfield and flourescent microscopy images of SLP-coated latex beads incubated with the indicated molecules diagramed at the top. Scale bars are given in the top left corner of each image and are 150μm. *See also Figure S4*.

In the context of engineering cSIgA variants that bind SLP, we also sought to introduce modifications that would allow us to distinguish between different IgA-based effector mechanisms such as coating, microscale crosslinking, classical agglutination, or enchained growth in experimental model systems. Having demonstrated that SC D2 can be replaced with a functional sdAb, we reasoned that other proteins or domains could replace D2 and confer detection such as fluorescence. Antibodies, typically IgGs, are frequently conjugated to fluorophores to facilitate fluorescence-based assays including fluorescence microscopy, Fluorescence Assisted Cell Sorting (FACS), Flow cytometry, and ELISA; however, the use of fluorescent secretory or polymeric IgA for these purposes and/or experiments in native mucosal environments is virtually non-existent. Accordingly, as a proof-of-principle we engineered cSC, in which SC D2 was replaced by a monomeric fluorescent proteins mCherry (**cSC^mCherry^**) or BFP (**cSC^BFP^)**.

Purified proteins were monodisperse and retained expected Absorbance and Fluorescence spectral properties (Figure 4A and Figure S4A). These data demonstrate that unliganded cSC can tolerate the replacement of D2 with an mFP, providing a means to enhance SIgA detection in vast mucosal environments.

With a goal of determining if cSC^mFP^ could be incorporated into liganded cSC (cSIgA), we created a library of cSC and cSIgA containing different combinations of antigen binding specificity and mCherry, all of which could be purified as monodisperse complexes. To test the ability of SLP-specific fluorescent, cS^mCherry^Fcα-CD5SLP, to bind SLP and confer fluorescence signal, we immobilized SLP on NHS-Agarose beads and visualized the beads in the presence or absence of cS^mCherry^Fcα-CD5SLP. Results demonstrated strong mCherry signal on SLP-Agarose beads in the presence of of cS^mCherry^Fcα-CD5SLP (Figure 4D). These results indicate that cSC^mFP^ can be effectively incorporated into cSIgA/cSFcα and used to support imaging and/or quantification of SIgA-antigen complexes, including the *C. difficile* SLP.

### cSIgA targeting SLP promotes C*. difficile* crosslinking

Having validated the functionality of our cSIgAs, we conducted 36-hour longitudinal experiments, in which *C. difficile* (ATCC-9869) cultures were inoculated in the presence or absence of antibodies, including sdCD5SLP, mFcα-CD5SLP, SFcα-CD5SLP, cS^mCherry^Fcα-CD5SLP and cS^CD5SLP^Fcα-CD5SLP and cS^mCherry^IgA-STA121 (non-binding control antibody). We monitored culture growth using optical density (OD_600nm_) (Figure 5A), qualitatively monitored cross-linking using confocal fluorescence microscopy and quantified crosslinking using Flow cytometry.

**Figure 5:**
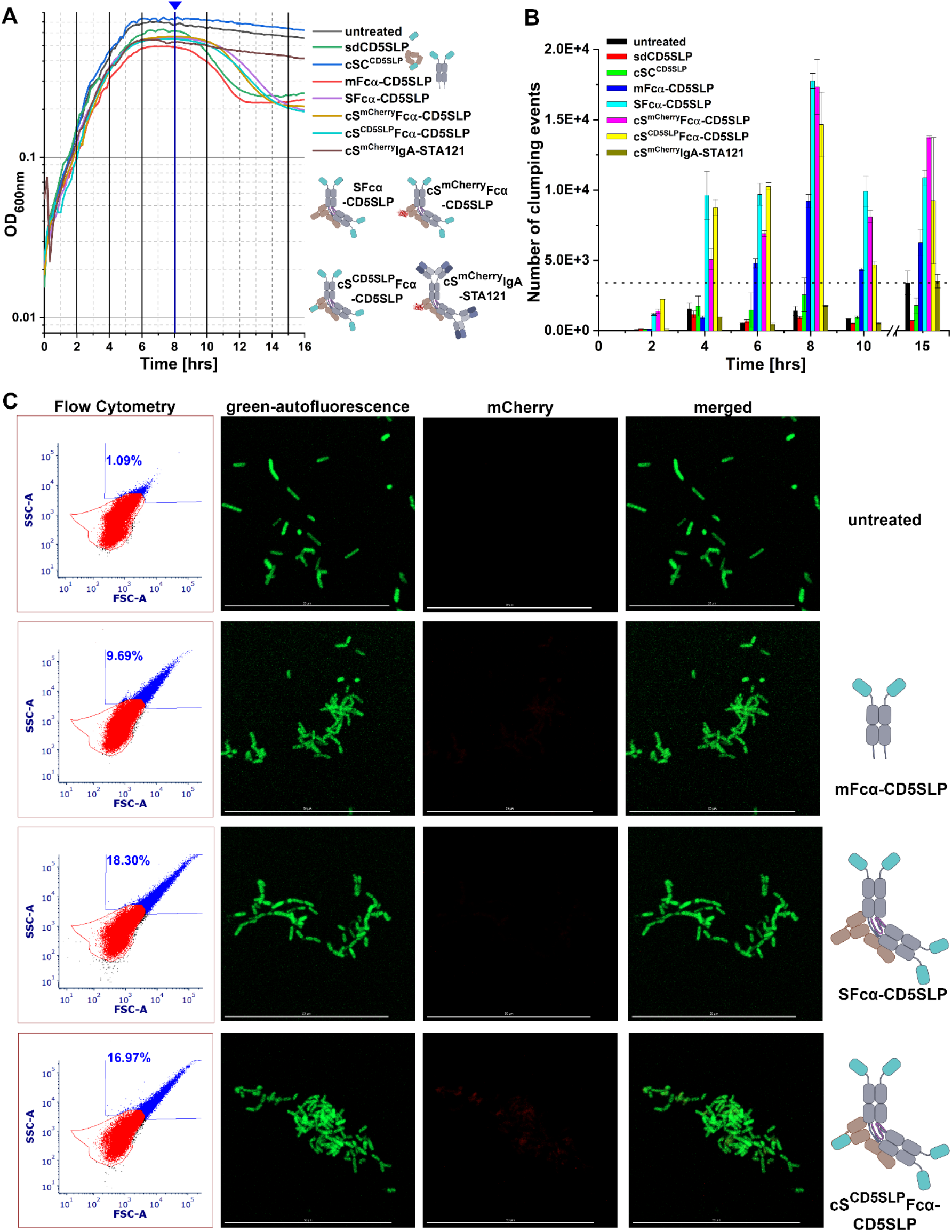
Polymeric anti-SLP antibodies cause *C. difficile* crosslinking. (A) OD_600nm_ versus time of the indicated samples shown in the key. The bold vertical grid lines indicate the time points at which samples were collected for Flow cytometry and imaging. The triangle denotes the timepoint at which number of clumping events is at its peak. (B) Shows the average number of clumping events recorded out of total 100,000 events for untreated *C. difficile* and those treated with the indicated antibody or control. (C) Flow cytometry analysis of *C. difficile* treated with the indicated antibodies or controls at the 8-hr. time point, and the respective confocal microscopy images. The scale bars are 50μm. *See also Figure S5 and Figure S6*.

Untreated *C. difficile*, cS^mCherry^IgA-STA121 (control), and cSC^CD5SLP^ treated *C. difficile* grew rapidly from the start, entering log phase after ∼1hrs, and reaching peak OD600 around ∼6hrs, followed by a steady decline of OD600 from ∼6 to 36hrs (Figure 5A). In contrast to untreated, cS^mCherry^IgA-STA121-treated or cSC^CD5SLP-^treated cultures, the sdCD5SLP, mFcα-CD5SLP, SFcα-CD5SLP, cS^mCherry^Fcα-CD5SLP, and cS^CD5SLP^Fcα-CD5SLP treated cultures exhibited rapid decline in OD600 following log and stationary phases (∼ 8-14 hours), after which OD600 remained constant (Figure 5A). It is noteworthy that unliganded cSC^CD5SLP^, showed similar growth trends to untreated and negative control antibody treated samples, likely indicating that *in-vitro* binding of a cSC^CD5SLP^ to the target antigen (Figure 5A) does not translate to notable effects on growing *C. difficile*.

Concurrently, we analyzed samples taken at 2hrs, 4hrs, 6hrs (log phase) and 8hrs, 10hrs, 15hrs (stationary phase and/or death phase) using Flow cytometry and confocal microscopy. The Flow cytometry data were optimized and gated with respect to the untreated *C. difficile* at 0hrs; the events within the gate were considered normal cell populations consisting of short chains of *C. difficile*. The events outside of this gate, corresponding to higher FSC-A and SSC-A, showed an increase in size and complexity and were categorized as clumped cells. We recorded the number of clumping events out of 100,000 events and analyzed changes in the value over time (Figure 5B).

Clumping was detected for *C. difficile* samples that were treated with chimeric antibodies containing at least two copies of CD5SLP, including mFcα-CD5SLP, SFcα-CD5SLP, cS^mCherry^Fcα-CD5SLP, and cS^CD5SLP^Fcα-CD5SLP. Untreated samples and those containing zero to one CD5SLP, cS^mCherry^IgA-STA121, cSC^CD5SLP^ or sdCD5SLP, showed a slow increase in the number of clumping events that plateaued around ∼3500 events (Figure 5B). This indicated that ∼3500 clumping events are expected for growing *C. difficile*, likely the result of dividing cells, crowding and non-specific interactions, and that clumping was not markedly impacted by the presence of antibodies with a single copy of CD5SLP. In contrast, mFcα-CD5SLP, SFcα-CD5SLP, cS^mCherry^Fcα-CD5SLP, and cS^CD5SLP^Fcα-CD5SLP exhibited 9217, 17764, 17312, 14656 clumping events for each sample, respectively, at the 8-hour time point, where maximum clumping was observed for most samples (Figure 5B). Notably, Flow cytometry data revealed a lag in clumping for mFcα-CD5SLP, which crosses the ∼3500 cutoff around 6hrs, in contrast to SFcα-CD5SLP, cS^mCherry^Fcα-CD5SLP, and cS^CD5SLP^Fcα-CD5SLP, which exceed 3500 events around 4hrs.

Confocal imaging of samples used for flow cytometry qualitatively demonstrated the clumping in cultures treated with antibodies having at least two CD5SLP, validating flow cytometry results (Figure 5C; Figure S5A). Together, data indicate that at least two copies of CD5SLP are needed to induce clumping but that clumping is enhanced when four or more copies of CD5SLP are included on one antibody. These results were consistent with replicate experiments that analyzed growth and quantified crosslinking using florescent microscopy images (Figure S6A to S6C). Furthermore, compared to confocal microscopy, crosslinking quantification using flow cytometry detected differences in clumping between antibodies containing two copies of CD5SLP versus four or five copies, that were not observed when counting cells in representative confocal images (Figure S6B). This may be in part due to using aliquots taken from culture samples, as opposed to the entire culture quantification via flow cytometry. Furthermore, we note that the size of clumps being detected via flow cytometry are likely larger than those designated as “clumping” in initial confocal microscopy experiments; in the host intestine, larger clumps of crosslinked bacteria are thought to promote removal via peristalsis. Notably, cS^mCherry^Fcα-CD5SLP exhibited clumping that was indistinguishable from other chimeras such as, mFcα-CD5SLP, SFcα-CD5SLP, cS^CD5SLP^Fcα-CD5SLP, indicating that the inclusion of mCherry does not measurably alter the process and could be used as a SIgA marker during experiments.

Inspection of confocal microscopy images corresponding to cS^mCherry^Fcα-CD5SLP-treated cultures at different time points revealed that *C. difficile* cells were coated with the antibody beginning at early time timepoints (2hrs). Between 2-6hrs (log-phase), images showed clumping of cells that appeared to be uniformly coated with cS^mCherry^Fcα-CD5SLP. However, around 8hrs (stationary phase), we observed non-uniform antibody coating on a subset of *C. difficile* cells within the culture, which correlated with morphological changes characterized by a “ruptured” appearance rather than the canonical rod-shaped structures (Figure 6). We also observed the ruptured morphology phenomenon in cultures treated with sdCD5SLP, mFcα-CD5SLP, SFcα-CD5SLP, and cS^CD5SLP^Fcα-CD5SLP between 8 to 36hrs to variable degrees, whereas cSC^CD5SLP^ showed minimal rupture only at 36hrs (Figure S7). The ruptured morphology was not observed for untreated and cS^mCherry^IgA-STA121-treated cultures. This indicates that antibodies targeting growing *C. difficile* SLP can induce morphological change in the microbe and that despite the lack of effector cells in our experiments, can have long-term disruptive effects on the pathogen.

**Figure 6:**
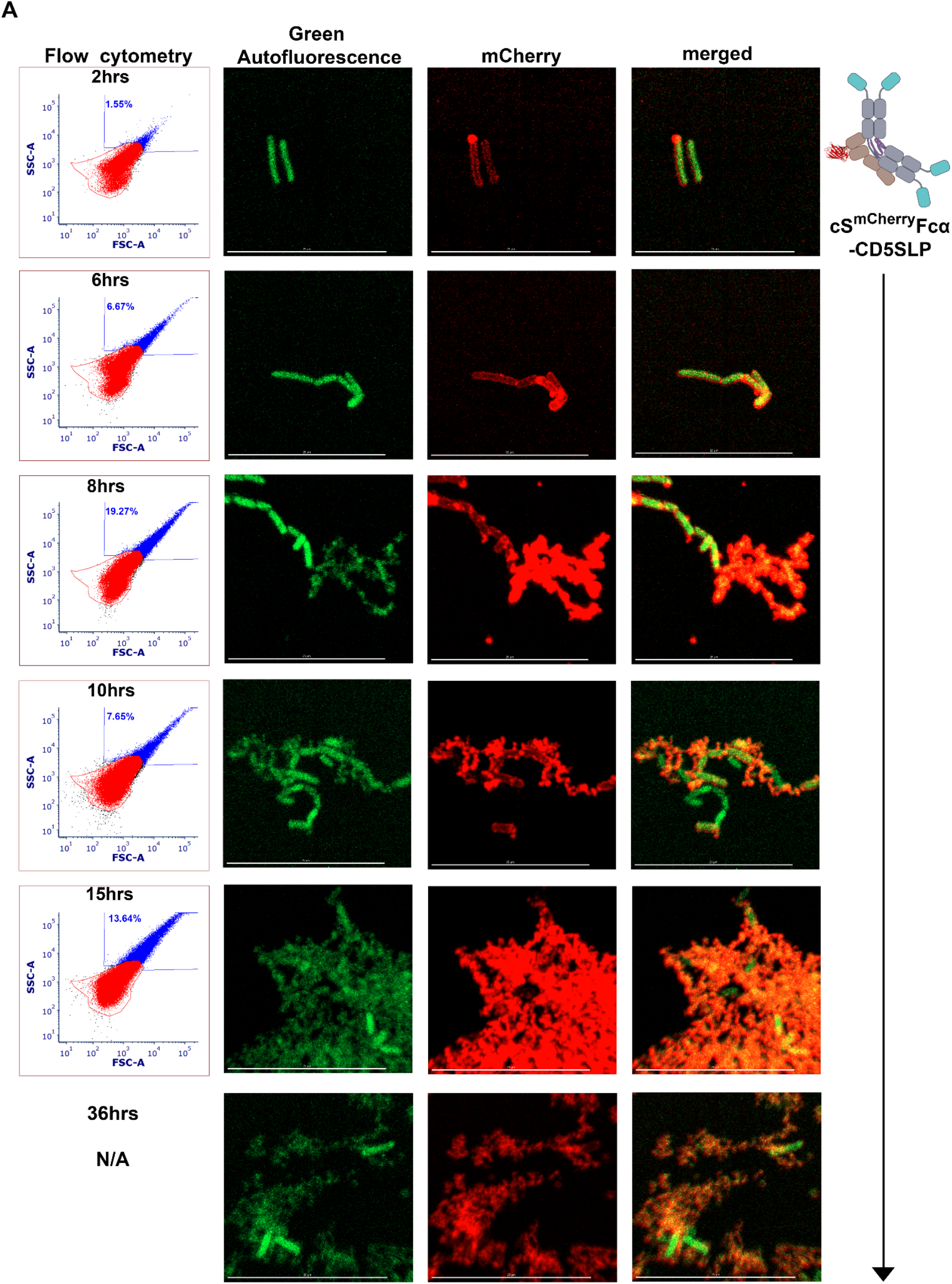
cS^mCherry^Fcα-CD5SLP coats *C. difficile* and causes morphological changes over time. Flow cytometry data and associated confocal fluorescent microscopy images from the indicated samples reveal the number of clumping events and the red fluorescence signal over time. The scales bars in all images are 25μm. *See also Figure S7*.

### Bi-specific cSIgA promotes *C. difficile* clumping and reduces cytotoxicity

Having determined that SIgAs targeting SLP can promote *C. difficile* clumping and having determined that cSIgAs can neutralize toxins, we engineered a bispecific cSIgA containing mFcα-CD5SLP and cSC^5D^ (TcdB targeting) called cS^5D^Fcα-CD5SLP. Consistent with prior experiments, cultures containing cS^5D^Fcα-CD5SLP exhibited clumping and eventual cell rupture (Figure 7A-C). Subsequently, we tested the cytotoxicity of supernatants from cultures treated with cS^5D^Fcα-CD5SLP or SFcα-CD5SLP, which lacks cSC^5D^ and encodes wild type SC, in Vero cell cytotoxicity assays. Compared to untreated controls, supernatants from cultures treated with cS^5D^Fcα-CD5SLP or SFcα-CD5SLP exhibited reduced cytotoxicity and cS^5D^Fcα-CD5SLP-treated cultures exhibited the greatest reduction in cytotoxicity (Figure 7B). These data suggest that the inclusion of CD5SLP and cSC^5D^ in a cSIgA can reduce TcdB toxicity in the culture microenvironment. While we anticipate that the cSC^5D^ reduces cytotoxicity directly as observed in experiments testing cSC^5D^ neutralization potency, the underlying mechanism for CD5SLP reduction in cytotoxicity is less clear. It may be due to an overall reduction in *C. difficile* growth in the presence of SLP-targeting antibodies, or due to changes in gene expression that lead to lower toxin production, or a combination of both. Regardless of the mechanism, SFcα-CD5SLP targeting SLPs appears to indirectly affect other virulence factors such as toxin production, suggesting that naturally occurring antibodies against SLP may help reduce pathogenicity and epithelial cell damage, in multiple ways. Together, results also highlight the therapeutic potential for SLP-targeting antibodies and bispecific antibodies to counter multiple aspects of pathogen virulence and minimize their cytotoxic effects.

**Figure 7:**
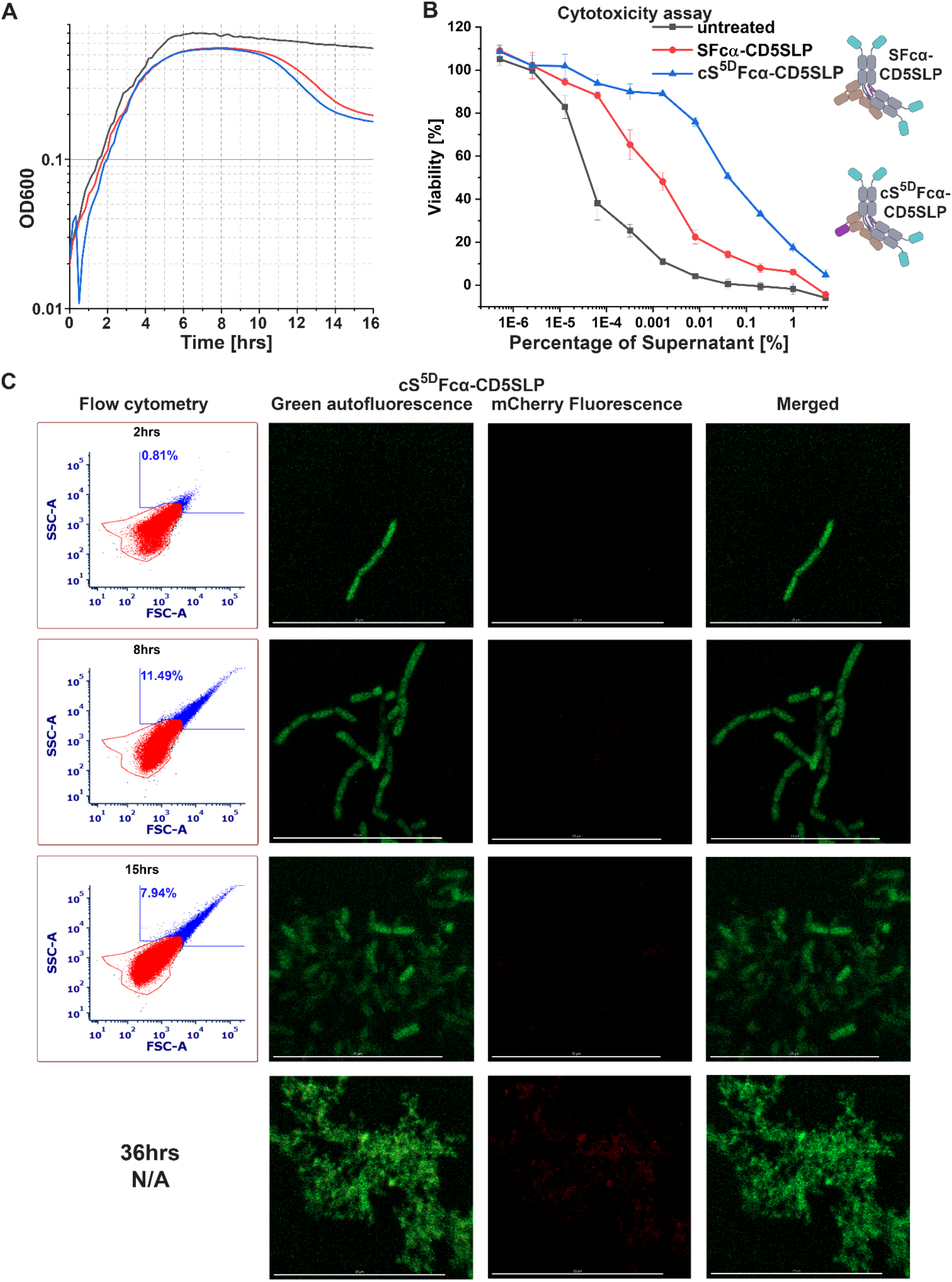
Bispecific antibody that targets *C. difficile* SLP and toxin, causes clumping as well as reduction is cytotoxicity. (A) Growth curves of *C. difficile*, growing in the presence of anti-SLP antibody SFcα-CD5SLP, and bispecific antibody cS^5D^Fcα-CD5SLP that targets SLP and TcdB. (B) The cytotoxicity of the supernatant obtained from cultures treated with the indicated antibodies. (C) Flow cytometry and fluorescent microscopy images of *C. difficile* cultures treated with cS^5D^Fcα-CD5SLP at the indicated time points. The scale bar in all images is 25μm.

## DISCUSSION

Humans devote considerable energy to SIgA production, yet the molecular mechanisms governing its unique effector mechanisms and impacts on diverse mucosal antigens remains poorly understood. Our work reveals that SIgA is surprisingly tolerant of large-scale structural modifications, which in turn have allowed us to explore its effector mechanisms and its developability as a therapeutic. We have focused on SIgA-dependent antigen neutralization and immune exclusion effector mechanisms, which are thought to dominate in mucosal secretions where immune effector cells and inflammation are limited[11],[47]. It is presumed that these mechanisms are supported by SIgA’s unique molecular structure, which cryo-EM structures and Fabs modeling revealed is conformationally asymmetric and likely to constrain the potential position Fabs can occupy, while leaving SC and two FcR binding sites sterically accessible[17],[30]. These geometric constraints are hypothesized to influence SIgA binding to antigens and other factors in unique ways, perhaps favoring interactions with some by promoting high avidity interactions and/or being optimal spaced to bind others[17]. It is often presumed that SIgA’s typically polymeric arrangement provides a crosslinking advantage over monomeric antibodies, or that SIgA will induce crosslinking (as opposed to coating) when bound to a given antigen; yet in practice, this has rarely been demonstrated or quantified experimentally, nor the impacts on host or microbe elucidated. For example, while humans develop SIgA’s against *C. difficile* SLP, whether binding to the antigen will result in coating, nano-scale crosslinking (between SLPs on one microbe), microbe-scale crosslinking, or some other effect and whether that SIgA provides an advantage over a monomeric antibody has remained unknown. Here we have tackled these and related questions, broadly demonstrating (1) that increasing the number of identical epitope binding sites on IgA can enhance soluble antigen neutralization (e.g. TcdA) (2) combining different epitope binding sites on a single IgA can promote neutralization of multiple soluble antigens (e.g TcdA and TcdB by cS^A20.1^IgA-PA41) (3) increasing the number of identical epitope binding sites on a single IgA can enhance immune-exclusion mechanisms, such as microbial crosslinking and/or target cell rupture (4) combining different epitope binding sites on a single cSIgA that targets multiple virulence factor epitopes (e.g. SLP and TcdB) can enhance immune exclusion and antigen neutralization and (5) combining epitope binding sites and mFP on a single cSIgA can facilitate visualization and quantification of cSIgA-antigen interactions and associated outcomes such as crosslinking. Together, these results demonstrate advantages of SIgA over monomeric antibodies applicable to understanding natural SIgA effector mechanisms, particularly those relevant to countering CDI, and demonstrate that SIgA’s beneficial attributes can be enhanced.

Our focus on SIgA interactions with *C. difficile* antigens stems from limited understanding of SIgA’s impact on CDI and the numerous challenges of using antibiotic treatments, including incomplete pathogen clearance, diseases recurrence, and the rise of antibiotic resistant human pathogens. These challenges have driven the development and FDA approval of Bezlotoxumab and signify that mAb treatments are a viable alternative treatment for CDI; however, Bezlotoxumab is administered intravenously[26], implying that it targets TcdB after epithelial barrier breach, whereas natural or engineered SIgAs have potential to target *C. difficile* antigens in the mucosa and limit infection before damage occurs. Indeed, our data demonstrate that in the case of toxin neutralization, increasing the number of binding sites is advantageous and that SIgA and cSIgA, containing four or five copies of sdA20.1 exhibited increased neutralization potency compared to variants containing one to two copies of sdA20.1. Unexpectedly, we found that cSC^A20.1^ could neutralize TcdA, but sdA20.1 alone did not. We speculate this advantage may arise from the larger, bulkier size of cSC^A20.1^, which may better block TcdA interactions with host cell receptors. sdA20.1 targets the TcdA CROP domain, however, CROPs are not the sole receptor binding region, and toxins lacking CROPs maintain some level of cytotoxicity[48],[49]. Therefore, cSC^A20.1^ may bind a CROP but also sterically occlude an adjacent receptor binding site. This finding is notable because while it appears that blocking one host receptor binding site (e.g. the sdA20.1 binding site) is an important part of the neutralization mechanism, it also hints that other factors are needed to achieve neutralization in the context of infection. Notably, colostrum SC has been implicated in limiting the effects of *C. difficile* toxins[18]; the innate-like mechanisms underlying this action is determined to be Glycan mediated[18]; however, we do not observe evidence that SC neutralizes toxins in our experiment, perhaps due to different glycosylation profiles of our recombinant SC or differences in experimental conditions, but when combined with sdA20.1, SC offers some specific advantage. We speculate that in the case of SIgA or cSIgA containing multiple copies of sdA20.1 (or another toxin-binding variable domain) not only would the size of SIgA help occlude receptor binding, but SIgA-dependent toxin crosslinking and the formation of multi-toxin soluble aggregates might further limit toxin access to host cell receptors by effectively blocking site not directly targeted by the antibody. Furthermore, in our experiments, we also observed synergistic effects when multiple toxins were targeted by a single cSIgA; the formation of multi-toxin aggregates and/or the ability of cSIgA binding at one toxin epitope to block receptor engagement of another may account for the synergistic effect we observe. In the case of *C. difficile* strains that express multiple toxins (such as hypervirulent *C. difficile* ribotypes BI/NAP1/027[50]), cSIgA may provide a treatment advantage.

*C. difficile* toxins are responsible for the vast majority of epithelial cell damage associated with CDI; however, persistent microbial growth is a major contributing factor for recurrence. Although humans develop antibodies against *C. difficile* surface antigens, whether these antibodies effectively exclude *C. difficile* from epithelial barriers, limit their growth and/or promote their expulsion has not been investigated[28]. Prior studies demonstrated that SIgAs targeting *Salmonella Typhimurium* (STm) surface antigens can promote immune exclusion via enchained growth and classical agglutination mechanisms and can also limit horizontal gene transfer and/or alter the surface of the microbe[16],[17],[51],[52]. Our experiments revealed that antibodies targeting SLP can influence cell viability and that those with two or more antigen binding sites, such as mFcα-CD5SLP, SFcα-CD5SLP, cS^mCherry^Fcα-CD5SLP, and cS^CD5SLP^Fcα-CD5SLP, can effectively induce crosslinking of *C. difficile* cells compared to the untreated controls, negative control antibodies, or those having a single SLP binding site, such as sdCD5SLP and cSC^CD5SLP^. Notably, those having four or more antigen binding sites exhibited even greater cross-linking potential. This indicates that in the context of a naturally occurring gut infection, SIgA targeting SLP has high potential to effectively limit *C. difficile* via immune exclusion mechanisms. Our data revealed that prior to crosslinking, SIgA coated bacterial cells and that subsequent crosslinking correlated with high cell density. This suggests that SIgA binding to SLP on the surface of one microbe does not prevent binding to the surface of another microbe and suggests that classical agglutination is a dominant mechanism. Classical agglutination is driven by the probability of cell-to-cell collisions, which increase with density, as opposed to enchained growth, which can occur at lower cell density when mother and daughter cells remain tethered. While we cannot rule out the possibility that enchained growth contributes to some percentage of the crosslinking we observe, we note that in addition to crosslinking occurring a high cell density, our images revealed clumped cells in many orientations, rather than long chains of cells connected end-to end, while untreated *C. difficile* cells are known to appear as rod-shaped and often in pairs or short chains[53]. In context of a naturally occurring infection and associated host antibody response, SIgA-SLP dependent agglutination would likely promote bacterial exclusion from epithelial cells and promote removal via peristalsis. Interestingly, our data also indicate that prolonged exposure to anti SLP antibodies can promote the rupture and death of *C. difficile* cells. The rupture of *C. difficile* appears to be independent of crosslinking since sdCD5SLP, and to lesser extent cSC^CD5SLP^, also caused cell rupture.

While our studies have provided new insights into SIgA effector mechanisms, they have also provided evidence that SIgA can be modified to enhance its beneficial properties. In the context of cSIgA targeting two soluble *C. difficile* toxins, TcdA and TcdB, we observed synergistic effects when compared to targeting each individually. Specifically, that cS^A20.1^IgA-PA41 showed enhanced neutralization of TcdA, as well as dual toxin neutralization of both TcdA and TcdB. We also found that we could combine TcdA and SLP-binding ability into a single SIgA that could both promote *C. difficile* cell crosslinking and reduce cytotoxicity. Surprisingly we also observe advantages of cSC over sdAbs, as evidenced by the ability of cSC^A20.1^ but not sdA20.1 to neutralize TcdA. Mammals secrete a large amount of free SC into the mucosa and while the reason for this remains uncertain, free SC may provide innate-like protection against some microbes, including *C. difficile* [18],[12]. SC is thought to be stable in mucosal secretions; it contains multiple N-linked glycans and adopts a compact closed structure[29], which we speculate may enhance the activity, stability and or localization (e.g. nonspecific binding) of the incorporated sdAb. We note, however, that SC may not always provide an advantage because cSC^CD5SLP^ failed to induce death phase or promote cell rupture compared to sdCD5SLP and thus, when targeting any antigen multiple factors will need to be tested to determine which form of antibody is most beneficial. Regardless, our data highlight the exciting possibility that engineered SIgA has potential to treat infections such as CDI. We envision that cSIgA biologics could be delivered to the site of infection (e.g. the large intestine or colon) to treat disease or prevent disease in high risk individuals.

Here we have focused efforts on a subset of SIgA effector mechanisms and targeted *C. difficile* antigens, but more broadly our results demonstrating that multiple sdAbs or mFPs can replace SC-D2 indicate that virtually any sdAb or mFP could replace SC-D2, providing modalities to track SC or SIgA and/or to enhance its effector mechanisms. This, combined with our ability to stably co-express cSC with different dIgAs illustrates the potential to generate diverse libraries of cSIgA that combine multiple antigen binding sites and/or different antigen binding sites and/or detection methods. For example, we co-expressed cSC^mCherry^ and cSC^mBFP^ dIgA to form cS^mFP^IgA, that we successfully utilized for antigen specific imaging and quantification by flow cytometry, without compromising the functions of the Fabs or altering the overall structure or conformational advantages of SIgA. We also note that cSIgAs are not expected to alter SIgA’s FcR binding sites and could in principle be used to study FcR-dependent mechanisms, which also remain poorly understood. We envision that cS^mFP^IgA, could be advantageous over SIgA-fluorescent dye conjugates, providing the ability to easily distinguish between native SIgA and introduced cSIgA in experiments involving mucosal administration of antibodies (e.g. to a mouse gut) where milligram quantities would likely be needed. We also envision that cS^mFP^IgA will simplify the quantification of SIgA coating or crosslinking of microbes (e.g., using flow cytometry or single molecule fluorescence), allowing us to build better understanding of SIgA effector functions and therapeutic potential.

## MATERIALS AND METHODS

### Construct design and cloning

The coding region of human JC (UniProt-P01591, native signal peptide sequence) was codon optimized, synthesized, and cloned into pD2610v1 (ATUM) analogous to mouse JC constructs[30]. The human IgA2m1 expression construct was created by fusing respective variable domain sequences along with the TPA signal peptide (residues MDAMKRGLCCVLLLCGAVFVSPAGA) to the constant human IgA2m1 HC (AAT74071.1)[55] and inserting them into pD2610v13 (ATUM). Briefly, VH region gene segments for different antibodies were selected, codon optimized, synthesized (IDT) and fused with the IgA2m1 constant heavy chain domains using Overlap-extension PCR[56] and resulting fusions were Electra cloned into pD2610v13 (ATUM). Fc-fused sdAb variants were created using an analogous method, in which the segment encoding the sdAb was fused to the IgA2m1 CH2-CH3 domains. His-tagged Fcα were generated by fusing the TPA signal peptide and HHHHHHGS sequence to IgA2m1 CH2-CH3 sequence using PCR; the resulting gene segment was cloned into pD2610v13 vector using Electra-cloning (Atum). Human light chain constructs were created using an analogous approach; in which VL domains were fused with CL domains of human IgLambda-1 (UniProt-P0DOX8) sequence, with signal peptide sequence (residues METDTLLLWVLLLWVPGSTG) and inserted into pD2610v13 vector by Electra cloning. The ectodomain region of human polymeric Immunoglobin receptor (pIgR) called Secretory Component (SC) (UniProt-P01833), was similarly synthesized, codon optimized and cloned into both pD2610v1 and pD2610v13 using Electra cloning. Chimeric SC (cSC) replacing D2 were generated by overlap-extension PCR, using codon optimized synthetic DNA (IDT) encoding the chimeric moiety and sequence overlapping with D1 and D3. The resulting gene segments were cloned into pD2610v1 or pD2610v13 vector using Electra cloning (Atum). Constructs encoding sdAbs were created by fusing an N-terminal signal peptide and C-terminal hexahistidine tag to the published VHH sequence. The sequence was codon optimized, synthesized and Electra cloned into the pD2610v13 vector. The different VH, VL and VHH are listed in Table 1.1 below.

**Table 1.1.**
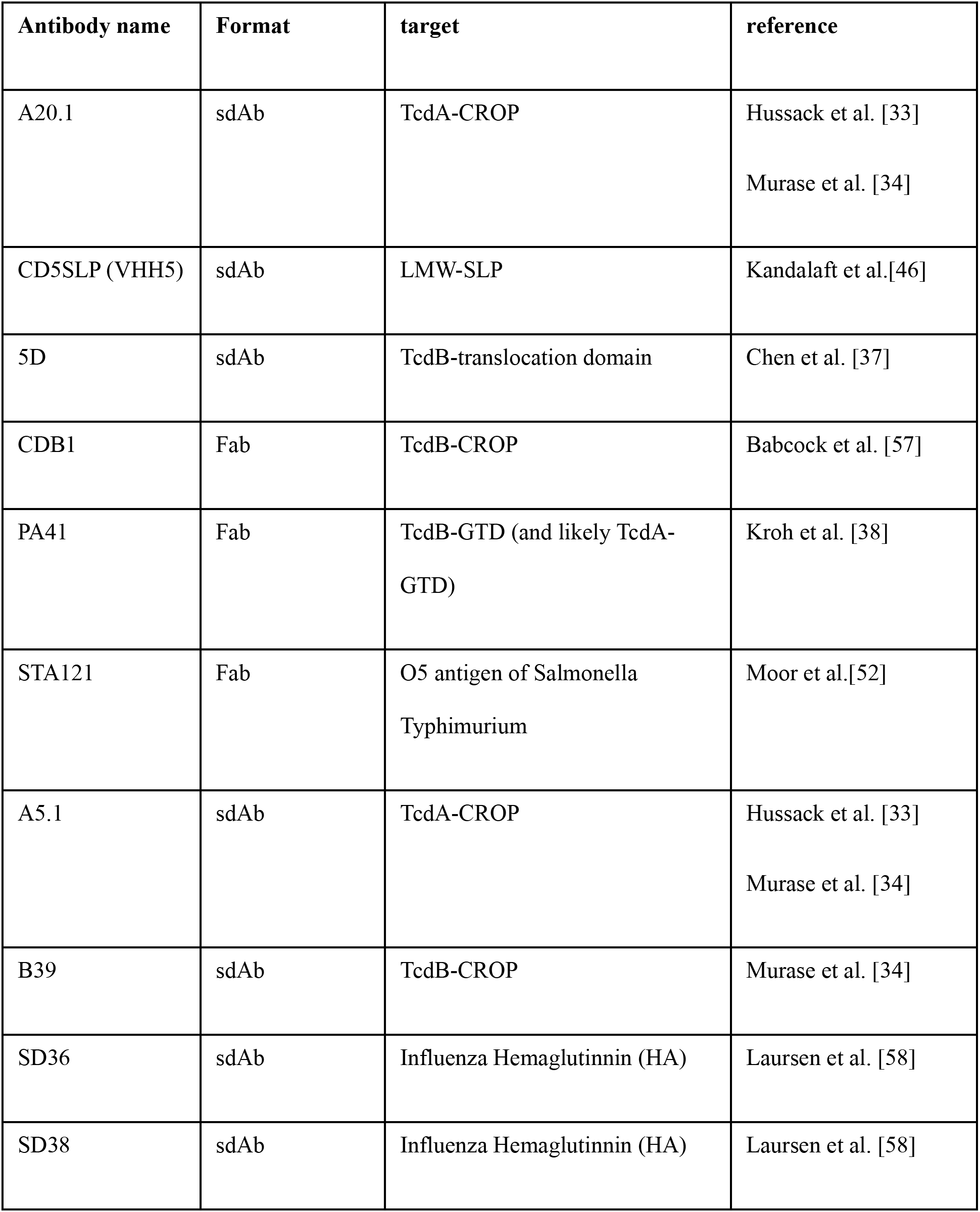
Antibody information.

### Transfection and harvesting

All constructs were transiently transfected (SC, cSC, sdAb, mFcα-fused-sdAb) or co-transfected (SFcα, SIgA, cSFcα, cSIgA) into Expi293F (Gibco: A14527) cells using the ExpiFectamine 293 transfection kit (ThermoFisher: A14525) and following previously described methods constructs[30]. Four to five days after transfection, supernatants were harvested and filtered through 0.22μm PES bottle top filters (Millipore Sigma).

### Bacterial protein expression

The plasmid for the expression of TXA1 (in pET28a vector) and SLP (in 2BCT vector) were provided by Dr Shannon Sirk’s Laboratory (UIUC). The plasmids were transformed into BL21 *E. coli* and screened with appropriate antibiotic resistance. The respective colonies were inoculated in 3ml starter culture, which was then used for larger 100-250ml culture to produce protein, by IPTG induction as per the New England Biolabs (NEB) protocol for BL21 (C2530). After 4 hours of protein expression at 37 degrees Celcius, the cells were pelleted at 4000xg. The pellets were resuspended in lysis buffer (50mM Tris 7.5, 500mM NaCl, 5-10% glycerol, 10mM Imidazole) containing lysozyme, DNAse and protease inhibitor cocktail (Pierce). The resuspended pellets were sonicated until the mixture had watery viscosity. The lysate was then centrifuged at 10000xg for 45 minutes, to remove cellular debris, and the supernatant was used for bacterial protein purification using Ni-NTA affinity resin (Qiagen: 30210).

### Protein Purification

The supernatants collected after transfection or the bacterial cell lysate, were used for purification of antibodies/ proteins with suitable affinity purification. SC, cSC, sdAb and proteins from bacterial cell lysates (SLP and TXA1) were purified with Ni-NTA resin (Qiagen: 30210), using Tris Buffered Saline (TBS: 20mM Tris+ 150mM NaCl, pH=7.4) + 250mM or 500ml imidazole for the elution. Similarly, His-Fcα constructs and complexes were purified by Ni-NTA purification methods and step elution gradient using TBS+250mM, TBS+500mM imidazole, TBS+750mM imidazole and TBS+1M imidazole. Fc-fused antibodies and full-length antibodies and their respective complexes with cSC were purified using Capture Select IgA affinity resin (Thermo Scientific: 194288010) and eluted using 0.1M glycine at pH=3, as per the manufacturer’s protocol. Following purification, all the proteins were buffer exchanged and maintained in 1X Phosphate Buffered Saline (PBS) pH=7.4 (Cytiva) using Amicon centrifugation concentrators (Millipore Sigma) with appropriate MW cutoff to buffer exchange and concentrate to a final volume of 0.5-1ml. The protein in PBS was 0.22um filtered using spin filters (Millipore Sigma) prior to Size exclusion Chromatography (SEC). SEC for the protein samples were carried out by using either Superose 6 column (Cytiva) or by using SD200 10 300 (Cytiva) column using AKTA pure (Cytiva) for automated equilibration, sample loading, and sample elution into 500ul elutants, and the respective absorbance data collection. The eluents after SEC were maintained in PBS until its use.

### Analytical SEC

Equi-molar ratios of protein-protein pairs were calculated and based on the estimated Molecular weight, which were determined from the ProtParam website (https://web.expasy.org/protparam). A minimum of 50ug of each protein was used for experiments. Based on calculations, equimolar amounts of the protein-protein mixture were combined in 1X PBS in final volume of 1ml, which was incubated at room temperature for 1hr and then 4°C before SEC. Controls consisted of the individual proteins in 1ml PBS at the same molarity as the mixture. The protein-protein mixture and the controls were subjected to SEC using SD200 10 300 column (Cytiva) attached to AKTA pure (Cytiva). The respective chromatograms were exported from the Unicorn 7.5 evaluation software (Cytiva) and plotted using OriginPro 2020 graphing and analysis software (OriginLab: https://www.originlab.com/).

### Fluorescence and absorbance measurements

For the chimeric fluorescent proteins, including cSC^mCherry^ or cSC^BFP^, the measurement of Fluorescence and the absorbance spectra was carried out using 1uM cSC in PBS in 100ul volume in a flat clear bottom 96 well plate (Thermo Scientific: 165305), and the absorbance and fluorescence spectra were measured using the Cytation 5 plate reader from BioTek. The absorbance was measured for appropriate range, and the Fluorescence was measured with 586nm excitation for cSC^mCherry^ and 400nm excitation for cSC^BFP^ and appropriate emission range. The respective absorbance and fluorescence spectra of PBS was used for background subtraction.

### NHS-SLP bead formation

Dry Pierce™ NHS-Activated Agarose and the purified Surface Layer Protein (SLP) in PBS were used (per the protocol for pierce NHS-activated agarose) for making SLP coated agarose beads. In a 1.5ml Eppendorf tube roughly 10-50ug (which translates to 75 to 375 ug of wet volume) of the NHS activated agarose was measured. To this, 1ml of 1uM Surface Layer Protein (SLP) in PBS was added to initiate the binding process. The binding was carried out by gently shaking at 4 degrees Celsius and 100 rpm. The NHS-agarose-SLP mixture was centrifuged at 1000xg and the supernatant removed, followed by two washes with 1ml PBS similarly by centrifuging at 1000xg. The final wet volume amounted to ∼250ug. To the washed agarose slurry 1M of ethanolamine (Sigma Aldrich: E9508) was added at 2 times the wet volume and incubated at room temperature for 20 minutes while shaking at 100rpm, in order to quench the binding of proteins and to block and inactivate the unoccupied NHS-agarose. This was followed by 3 washes with TBS and the final SLP-Agarose were stored in final volume of 1ml in TBS for experiments (250ug/ml agarose concentration approximately).

### Bead Fluorescence study

For the bead binding and imaging assays, SLP-conjugated bead slurry was agitated and 100ul of slurry was aliquoted into a1.5ml labelled Eppendorf tubes. The slurry was then centrifuged at 1000xg, and the supernatant was discarded and replaced with 100ul of 5uM analyte protein in TBS. The analyte-slurry mixture was allowed to incubate at room temperature for 1hr, following by three TBS washes, and final resuspension in 100ul TBS. 10-20ul of this final slurry, was placed on a plain glass slide (Fisher Scientific) with coverslip (Fisher Scientific) sealed with clear nail polish on the edges. The agarose beads were imaged using Leica inverted fluorescence microscope for brightfield and fluorescence imaging (Dr. Kai Zhang Lab). The images collected were visualized from Imaris Viewer software.

### Vero cell cytotoxicity neutralization assays

Vero cells (ATCC-CCL-81) were seeded in tissue culture treated 96 well plates (Thermo Scientific), at 10000cells/well in complete media, which is Dulbecco’s Modified Eagle Medium (DMEM) (Gibco) + 10% Fetal Bovine Serum (FBS) (Gibco), and then incubated in 8% CO2 at 37deg Celsius (day 0). After 20-22hrs (day1), the complete media from each well was removed, and the preincubated mixture of respective concentrations of toxins and antibodies in complete media were added to the respective 96 wells. The plates were then incubated for 65hrs. Prior to measuring the cell viability the cells were washed with MEM (Earles MEM from Gibco) at least two times, and then a 1:10 mixture of Alamar blue cell viability reagent (Invitrogen: DAL1100) were prepared in MEM and 100ul of this mixture added to all the wells[36]. The plates were then incubated at 37degrees for 3hrs, and the fluorescence intensity was measured for each well at 560nm absorbance and 590nm emission using Cytation 5 Plate reader from Biotek. For TcdA assays, a 50pM concentration of TcdA was used; for TcdB assays 4pM of TcdB was used, and for dual toxin neutralization assays 50pM TcdA and 4pM TcdB were used. The antibody concentrations and its dilution series varied depending on the experimental requirements to get appropriate range of Vero cell viabilities for comparison within each experiment.

### *C. difficile* neutralization assays

*Clostridiodes difficile* ATCC-9689-fz, strain designation-90556-M6S, toxinotype 0 and ribotype 001, was used for all experiments. As per the protocol [59], [60], the *C. difficile* was optimized to grow in an anaerobic chamber (Coy Laboratories) at 37degrees Celcius. Brain Heart Infusion medium (37g/L) with yeast extract (5g/L) (BHIS) supplemented with 0.1% L-cysteine (Sigma Aldrich: 168149) were prepared, aliquoted into sterile culture tubes and pre-reduced in the anaerobic chamber for 1 day prior to *C. difficile* inoculation. Additionally, plastic products such as flat and clear bottom 96-well plates (Fisher Scientific), tips and other consumables were pre-reduced in the anaerobic chamber for at least 1 day. The *C. difficile* was inoculated from a frozen glycerol stock according to the published protocol [59], [60]. This 12-18hr overnight culture was used to set up *C. difficile* neutralization assays. The OD_600nm_ of the overnight was measured, and accordingly the samples were diluted to an estimated OD of 0.05 with BHIS+0.1% L-cysteine. In the 96 well plates, diluted *C. difficile* was mixed with equal volume of 8uM concentrations of antibodies diluted in BHIS, in 1:1 ratio to get final concentration of 4uM of antibody in the assay. The plates were sealed with breathable membrane (Millipore Sigma) to prevent condensation overtime and minimize evaporation, and Cerillo Stratus portable plate reader was used to measure OD_600nm_ every 10 mins. Duplicates plates were set up for sample extraction for Flow cytometry and Confocal microscopy slide preparation.

### Cytotoxicity of *C. difficile* growth supernatant

Cytotoxicity of *C. difficile* supernatant was measured by collecting supernatant samples at indicated OD_600nm_ measurements in the 96-well plates specific for the Cerillo Stratus plate reader. Briefly, this was carried out by spinning down the cells, and then filtering the supernatant through a 0.22um filter pior to collection in the plate. The cytotoxicity of supernatant for treated and untreated samples were analyzed using Vero Cells (ATCC-CCL-81) cultured as previously described. Cytotoxicity evaluations were carried out by administering 2-5% supernatant starting concentration and the respective 5-fold dilution series in DMEM+10%FBS to the pre-seeded 10000cells/well Vero cells. The cell viability was analyzed after 48 hours using the alamar blue assay as previously described. Appropriate positive control (complete media + 100pMTcdA + 10pM TcdB) and negative control (complete media) were used to estimate the viability.

### Confocal microscopy of *C. difficile* samples

At different time points during the growth for *C. difficile* in the presence and absence of antibodies, 5ul of samples were collected from the respective 96-well plates of *C. difficile* culture. 5ul samples were mixed with 5ul of 8% Paraformaldehyde (PFA) (Fisher Scientific) in PBS, and then placed on a glass slide (Fisher Scientific), and a coverslip (Fisher Scientific) was slowly placed using forceps and the edges of the sealed using nail polish. All the slide preps were carried out inside the anaerobic chamber. The wet mount microscopic slides were imaged using a Zeiss LSM 900, with 63X/oil and 1.4NA, at 1X zoom. The autofluorescence of *C. difficile* was highest in the green channel, so an excitation of 488nm and 0.5% of the laser intensity was used to image the green autofluorescence; for mCherry an excitation of 561nm and 0.5 %-2% laser intensity was used. The transmission photomultiplier tube (T-PMT) was set with 561nm in the same channel as mCherry for phase contrast data collection. Appropriate gains were used for green channel and red channels, optimized based on the samples with weakest fluorescence and the brightest fluorescence. Images were collected as tiles for manually counting, and/or with Z-stack as needed.

### Analyses of images

Bacterial clumping was quantified by manual counting of bacterial on the phase contrast images, using the Image-J cell counter plugin (https://imagej.nih.gov/ij/). The counting was carried out based on a defined set of rules: Clumped cells were defined as 3 or more cells in contact with each other in a non-end-to-end (non-linear) manner, and un-clumped cells were defined as cells which were single and dispersed and/or growing and connected in an end-to-end (linear) manner. The images for the figures were created using Imaris viewer (https://imaris.oxinst.com/imaris-viewer).

### Flow cytometry

100ul of untreated or antibody-treated *C. difficile* cultures were extracted from a 96-well culture plate, mixed with PFA to final concentration of 2% PFA and incubated at RT for at least 30 minutes. Fixed *C. difficile* cultures were diluted with PBS to a minimum volume of 400ul and loaded into a LSR Fortessa Flow cytometer. PMT voltages and thresholds were adjusted based on untreated *C. difficile* and the FITC and Texas red fluorophore setting was utilized to record the green and red fluorescence. 300,000 events were recorded for untreated *C. difficile* at the time point of 0 hours. All the other time points for treated and untreated samples, and their replicates, were recorded for 100,000 events. Samples taken at 20 hours and 36 hours were not analyzed by Flow cytometry due to predominant cell rupture.

### Flow cytometry data analysis

FCS Express 6 and 7, were utilized for Flow cytometry data analysis. All samples were gated in SSC-A vs FSC-A dot plot according to untreated *C. difficile* sample at T=0 hours of the experiment. The gate enclosing of >99% of *C. difficile* cells at T=0 hours were classified as the “normal population”, and another polygonal gate drawn outside this gate to account for higher SSC-A vs FSC-A were categorized as the “clumped population”. The respective statistical data of number of total clumped population were extracted from FCS Express and was plotted against time for each samples using OriginPro 2020. The dot-plots of SSC-A vs FSC-A were exported as images from FCS Express.

## ACKNOWLEDGEMENTS

We thank members of the Stadtmueller Laboratory for insightful conversations and suggestions related to this work. We thank Kritika Mehta, Dr. Kai Zhang, Dr. Elizabeth Roland, Dr. Collin Kieffer and the UIUC Carl R. Woese Institute for genomic Biology Imaging Core for microscopy support. We thank Michael Miller’s Lab (UIUC), especially Dr. Miller and Zifan Xie for providing laboratory space and training to conduct *C. difficile* experiments. We thank Dr. Craig Ellermeier (U. Iowa) for insightful discussions on *C. difficile* and Dr. Shannon Sirk for providing plasmids for *C. difficile* antigens (TXA1 and SLP). We also thankthe UIUC Roy. J. Carver Biotechnology center and the Cytometry and Microscopy to Omics (CMtO) Facility for the use of the BD LSR Fortessa for flow cytometry as well as the UIUC School of Chemical Sciences High Throughput Screening facility for use of their Biotek cytation 5 plate reader. This work was supported by NIH grant 1R01AI165570 and University of Illinois start-up funding to B.M.S.

## AUTHOR CONTRIBUTIONS

The study was conceived by B.M.S and S.K.B.; experiments were conducted by S.K.B; MJM provided laboratory space and critical advice supporting *C. difficile* experiments. All authors contributed to data analysis and manuscript writing.

## COMPETING INTERESTS

BMS and SKB are listed as inventors on a patent related to SIgA engineering (WO2023044419) [54]. (https://patentscope.wipo.int/search/en/detail.jsf?docId=WO2023044419&_cid=P12-LHI6X3-04314-1)

## Supplementary Figures

**Figure S1.**
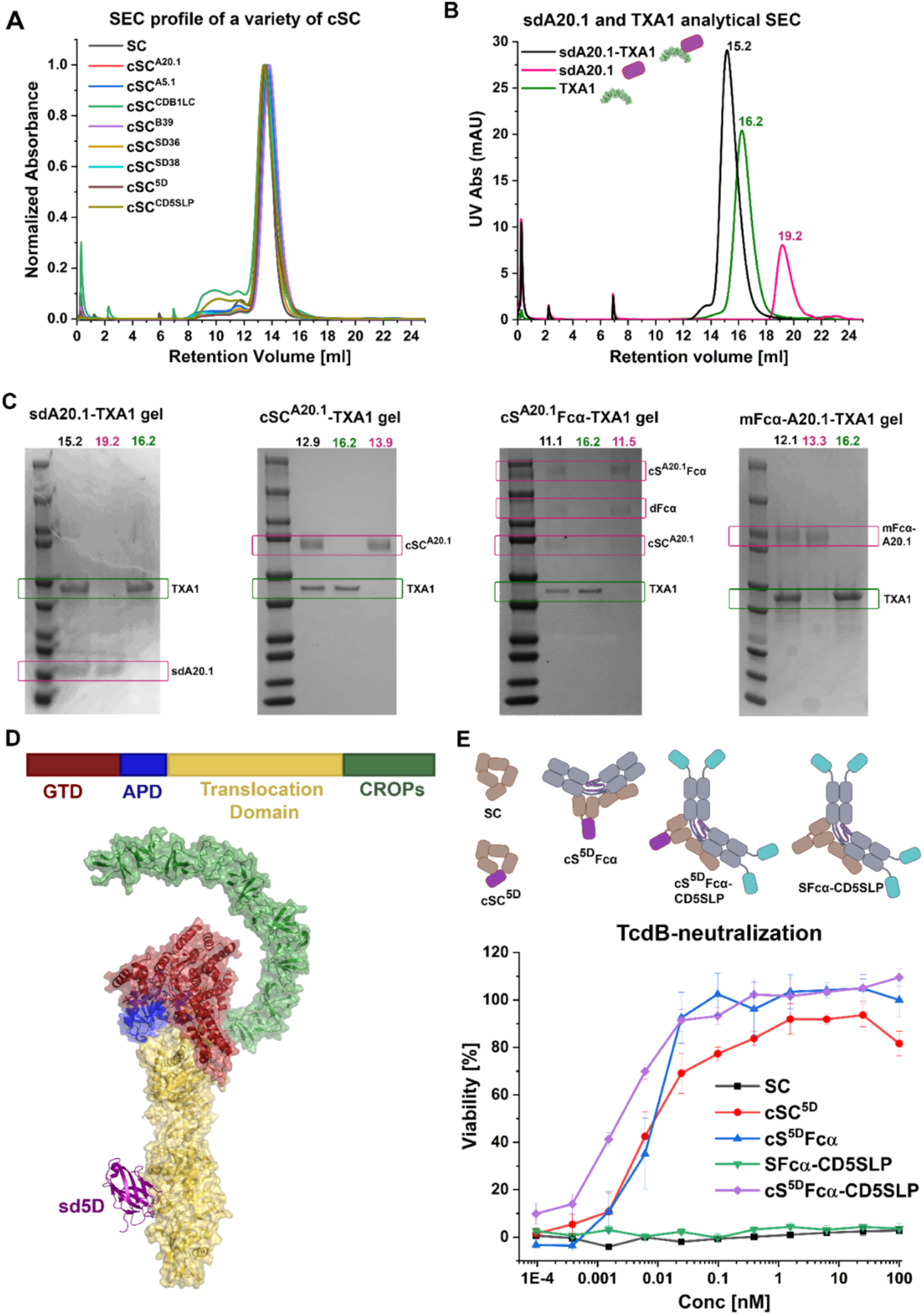
related to Figure 2: Functional cSC can be generated and utilized in its liganded and unliganded forms. (A) Normalized monodisperse SEC peaks of the indicated cSC variants, cSC^A20.1^, cSC^A5.1^, cSC^B39^, cSC^5D^, cSC^CDB1LC^. (B) Schematic and analytical SEC elution profiles of sdA20.1, TcdA fragment, TXA1, and the sdA20.1-TXA1 complex. The analytical SEC was carried out by incubation of equimolar amounts of TXA1 and sdA20.1 at room temperature for 1hour, followed by chilling the mixture to 4°C and then carrying out SEC. TXA1 and sdA20.1 were individually incubated similarly and subjected to SEC for comparison. (C) The SDS PAGE analysis of samples collected from the peaks of the analytical SEC samples. (D) Schematic of TcdB domain organization (top), and TcdB structure binding to sd5D (iodine purple) binding to the translocation domain (bottom, PDB:6OQ5). (E) Vero cell-based cytotoxicity neutralization assay showing the concentration dependent neutralization of 4pM TcdB by the indicated molecules including cSC^5D^ in liganded and unliganded. The schematics of all the antibodies used are shown on top.

**Figure S2.**
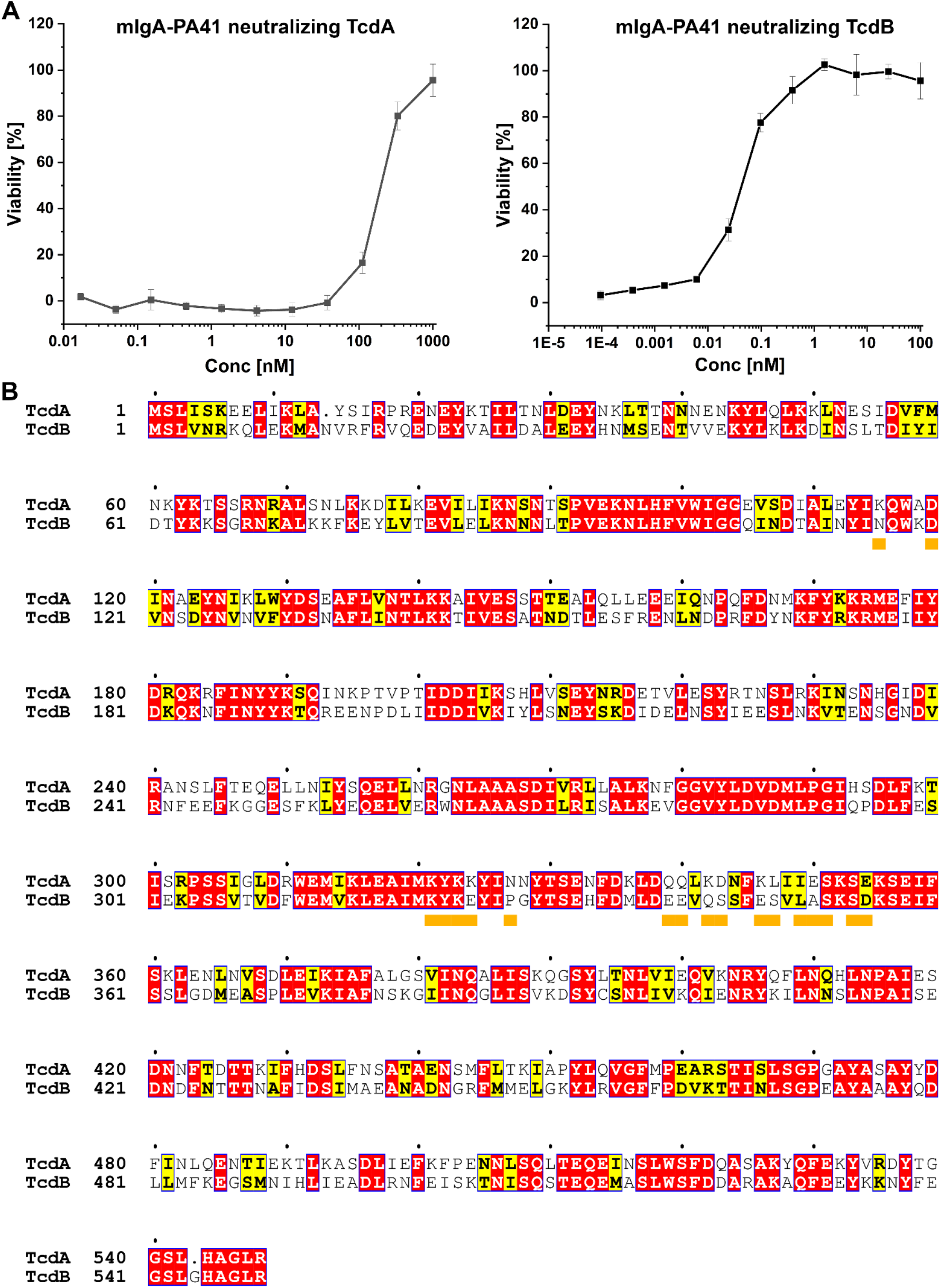
related to Figure 3 and Figure S3: The PA41 antibody neutralizes both TcdA and TcdB. (A) mIgA-PA41 neutralization of TcdA (50pM) and TcdB (4pM). (B) Sequence alignment of the Glucosyl Transferase domain (GTD) from TcdA and TcdB; the contacting residues of TcdB with PA41 are indicated by orange squares.

**Figure S3.**
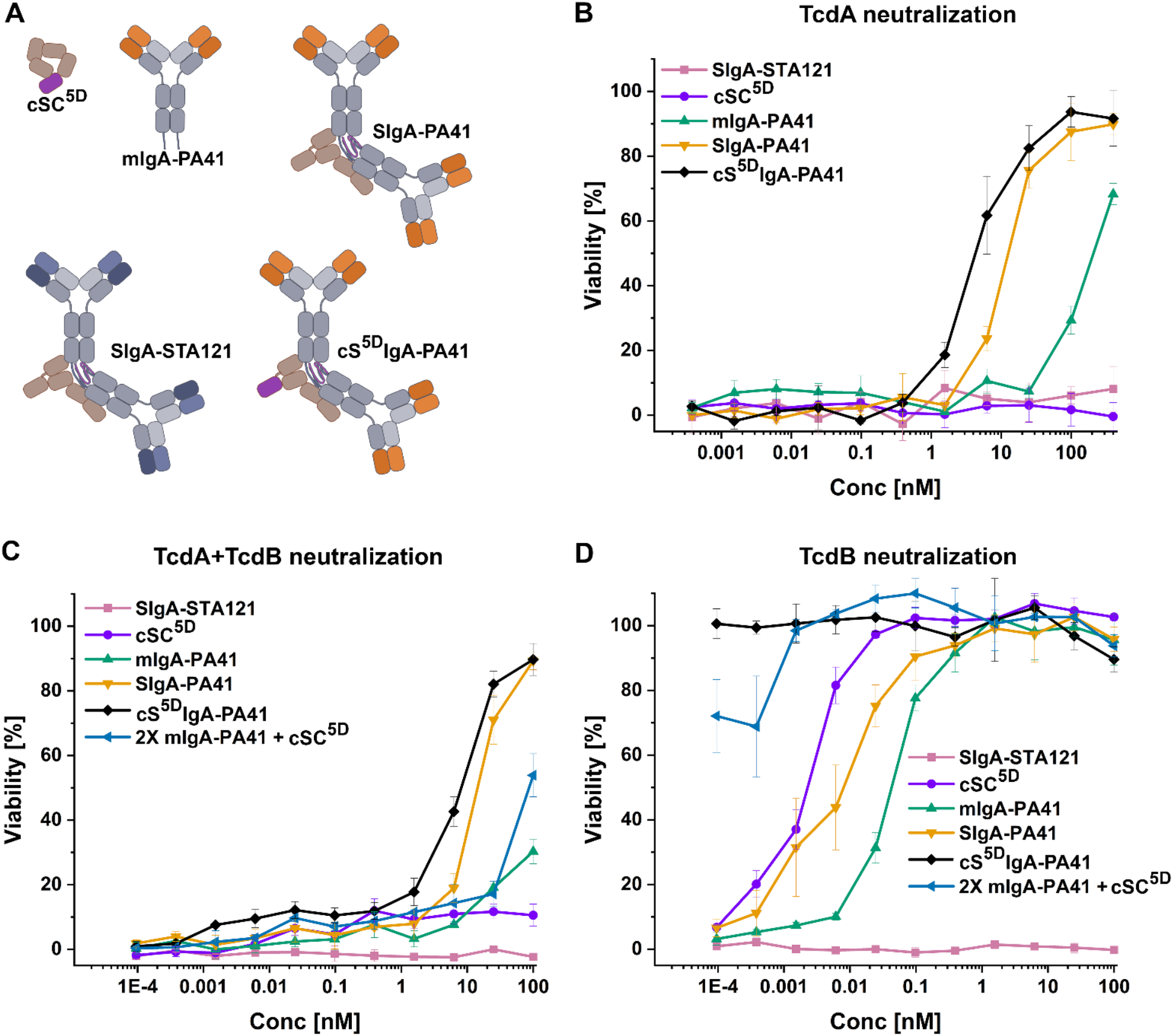
related to Figure 3 and Figure S2: The synergistic neutralization effects of bispecific cS^5D^IgA-PA41. (A) Schematic of antibodies used for the Vero cell-based cytotoxicity neutralization assay specifically to test cS^5D^IgA-PA41, where the sd5D moiety neutralizes TcdB, and PA41-Fab neutralizes TcdA and TcdB. (B) Concentration-based neutralization TcdA (50pM) by the indicated antibodies. (C) Concentration dependent neutralization of TcdA (50pM) and TcdB (4pM) by the indicated antibodies. (D) Concentration dependent neutralization of TcdB (4pM) by the indicated antibodies.

**Figure S4.**
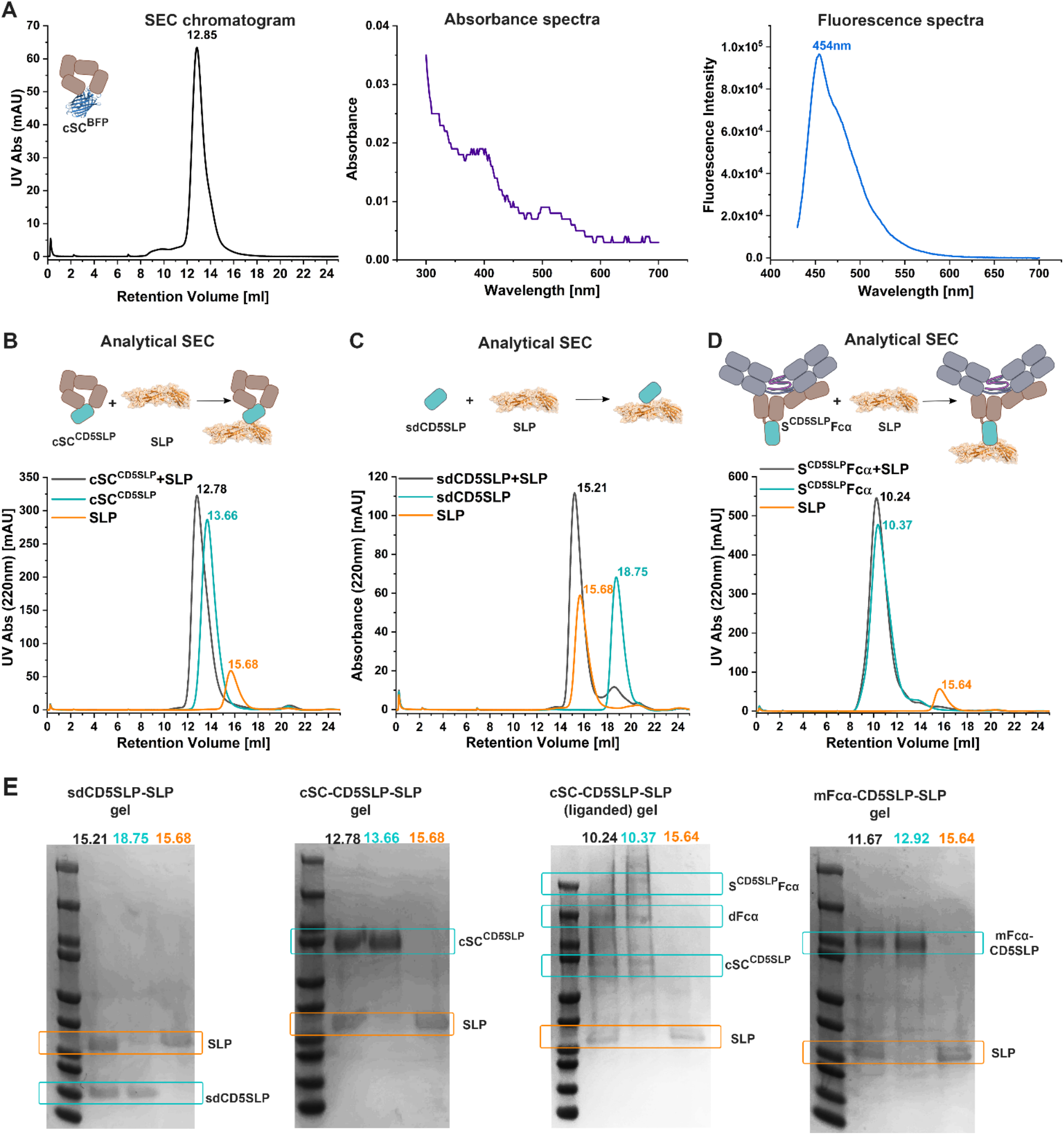
related to Figure 4. (A) The SEC chromatogram, absorbance spectra and fluorescence spectra of mSC^BFP^. (B) Analytical SEC showing the binding of cSC^CD5SLP^ and SLP. (C) Analytical SEC showing the binding of sdCD5SLP and SLP. (D) Analytical SEC showing the binding of S^CD5SLP^Fcα (liganded cSC^CD5SLP^) and SLP. (E) The SDS PAGE analysis of samples collected from the peaks of the analytical SEC samples.

**Figure S5.**
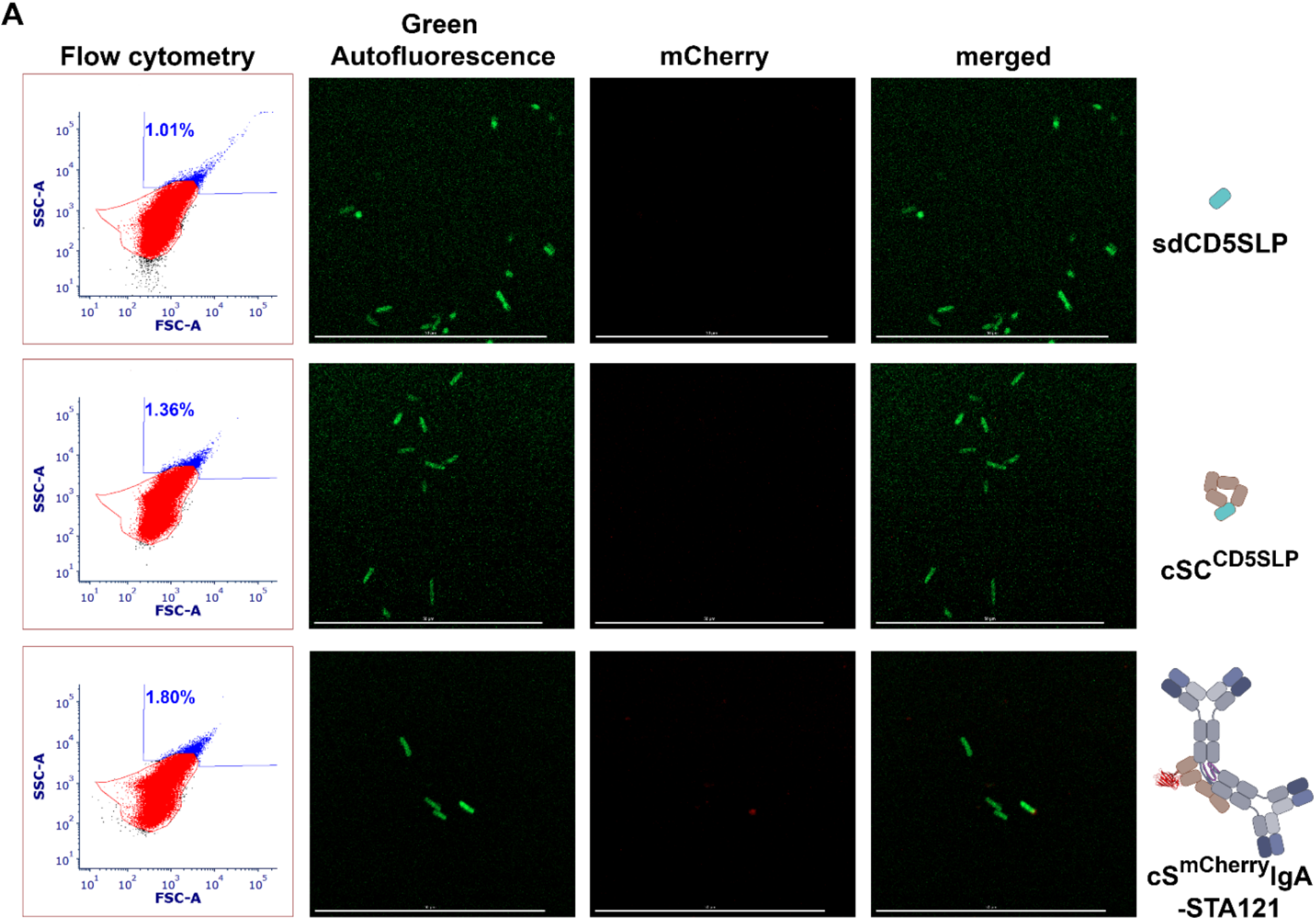
related to Figure 5: Antibodies with one SLP binding sites do not promote crosslinking. (A) Flow cytometry data for *C. difficile* samples incubated with the indicated antibody or control at the 8-hr. time point, and the associated confocal microscopy images. The scale bars are 50μm.

**Figure S6.**
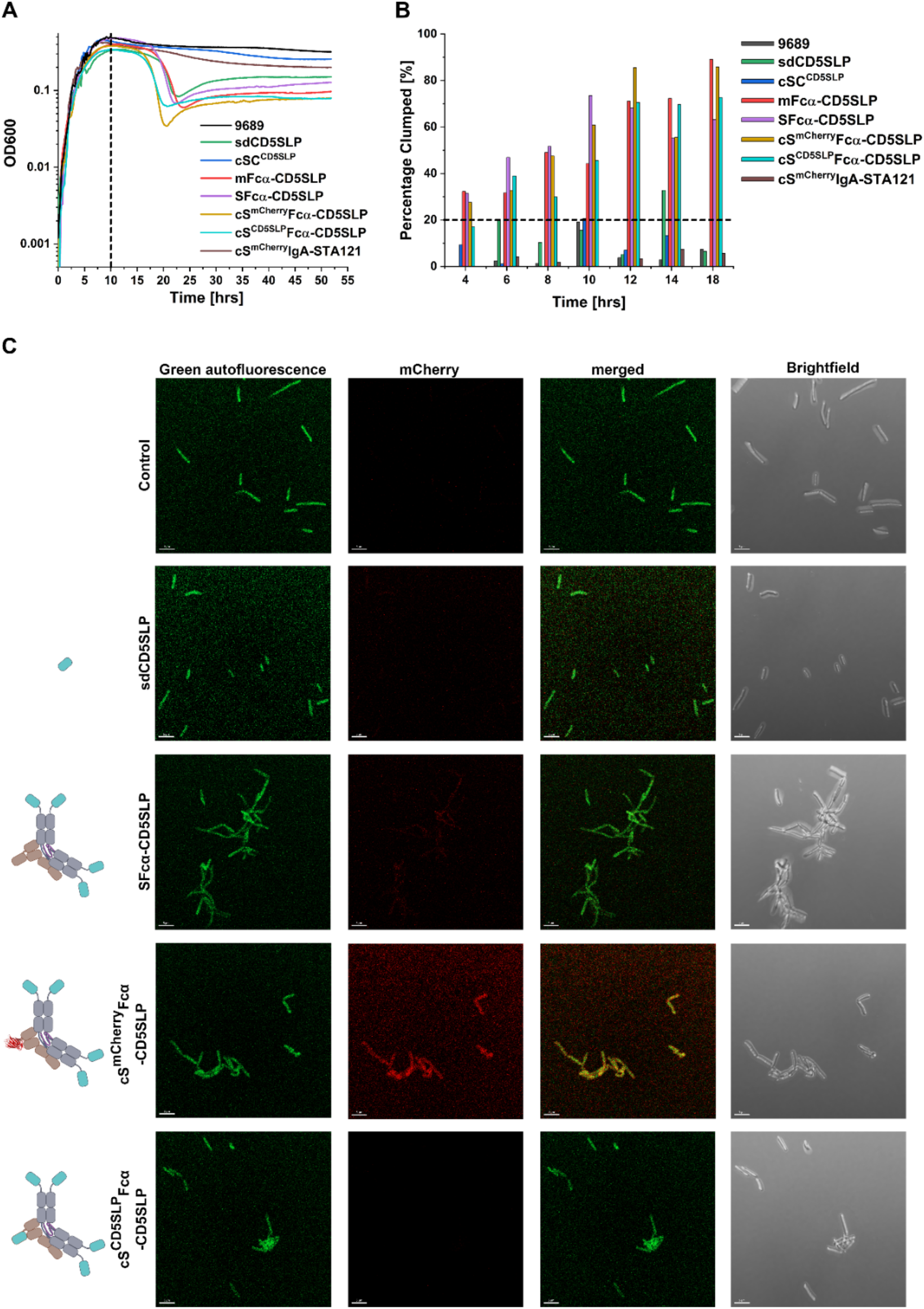
related to Figure 5: State of *C. difficile* clumping for different samples at the peak of growth phase. (A) Replicate experiment growth curves for the indicated antibodies or controls administered to growing *C. difficile* cultures. (B) The percentage of clumped cells in each of the indicated samples as determined by manual counting of >100 cells/sample (error bars statistically not applicable for heterogenous sample, and variables). (C) Representative confocal and bright field microscopy images for the indicated antibodies administered to *C. difficile* at the 10-hour time point in the experiment. The scale bars are 5μm.

**Figure S7.**
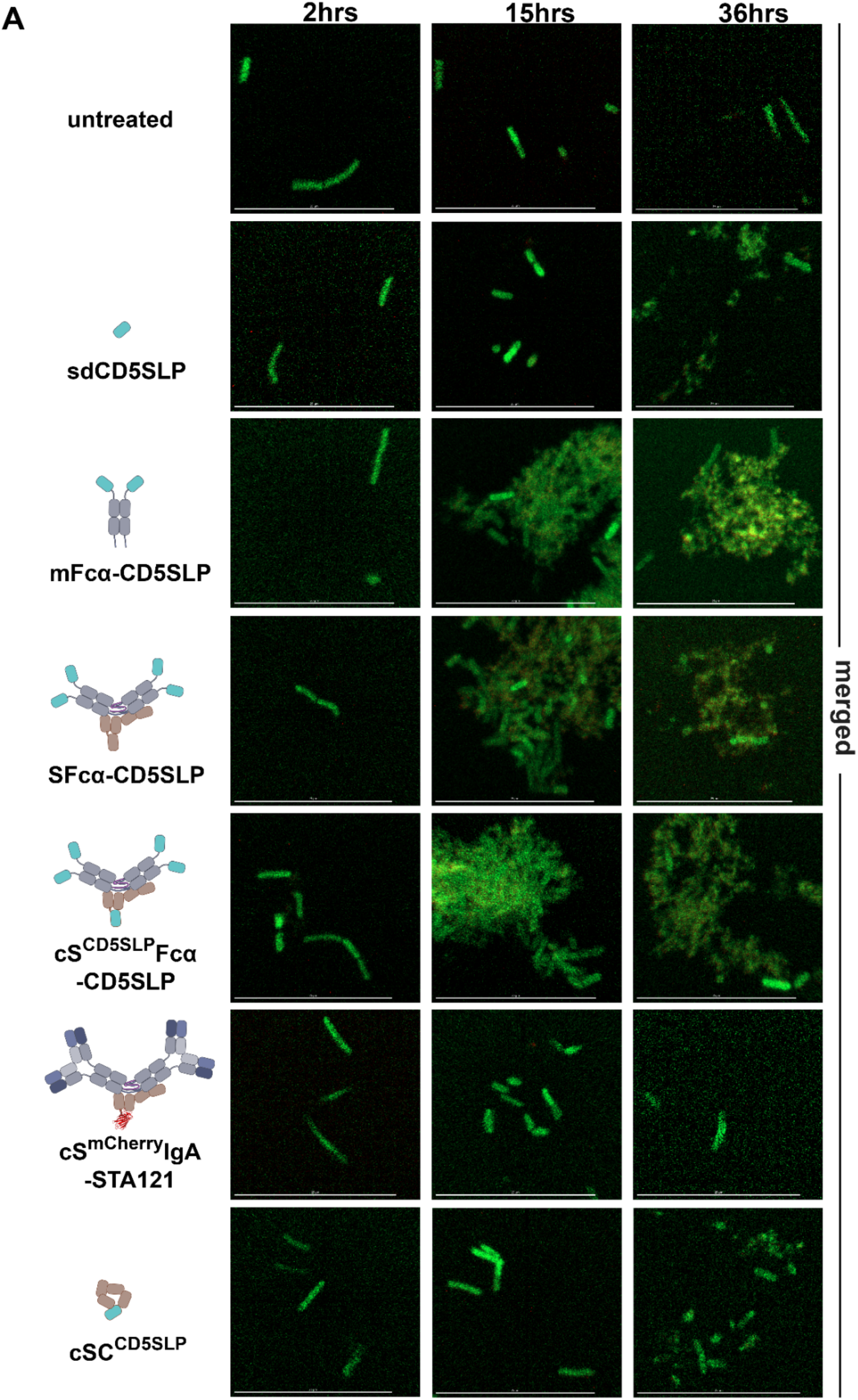
related to Figure 6: State of *C. difficile clumping at* three time points. (A) Confocal microscopy images detailing *C. difficile* cultures in the indicated samples. The scales scale bars in all images are 25μm.

## Notes

### Competing Interest Statement

Authors BMS and SKB are listed as inventors on patent WO2023044419-CHIMERIC SECRETORY COMPONENT POLYPEPTIDES AND USES THEREOF.

